# Developmental role of macrophages modelled in human pluripotent stem cell derived intestinal tissue

**DOI:** 10.1101/2022.09.09.505715

**Authors:** Andrew T. Song, Renata H. M. Sindeaux, Yuanyi Li, Hicham Affia, Tapan Agnihotri, Severine Leclerc, Patrick Piet van Vliet, Mathieu Colas, Jean-Victor Guimond, Natasha Patey, Jean-Sebastien Joyal, Elie Haddad, Luis Barreiro, Gregor Andelfinger

## Abstract

Macrophages populate the embryo early in gestation but their role in the developmental process remains largely unknown. In particular, specification and function of macrophages in intestinal development remain unexplored. To study this event in human developmental context, we derived and combined human intestinal organoid and macrophages from pluripotent stem cells. Macrophages migrated into the organoid, proliferated, and occupied the emerging micro-anatomical niches of epithelial crypts and ganglia. They also acquired a similar transcriptomic profile to fetal intestinal macrophages and displayed tissue macrophage behaviors, such as recruitment to tissue injury. Using this model, we show that macrophages reduce glycolysis in mesenchymal cells and limit tissue growth without affecting tissue architecture, in contrast to the pro-growth effect of enteric neurons. In short, we engineered an intestinal tissue model populated with macrophages, and we suggest that resident macrophages contribute to regulation of metabolism and growth of the developing intestine.

## Introduction

Macrophages arise early in the embryonic development, spread throughout the organism, and adopt site-specific fates required to maintain normal tissue functions^1^. The intestine, for instance, harbors a range of macrophage subtypes in distinct locations. Within the villi, they phagocytose contents of the lumen through the epithelium^2–4^. At the base of the epithelium, crypt associated macrophages regulate epithelial stemness, and differentiation^5–7^. The enteric nervous system collaborates with adjacent macrophages to regulate immune response and peristalsis^8–10^. In contrast to our growing knowledge in intestinal macrophages in adult organisms, it remains unknown if they influence the developmental process itself.

Accumulating evidence in mice suggests that macrophages affect tissue development. In embryonic development, they have been shown to regulate growth of kidney, development of pancreatic islet, and formation of coronary plexus and lymphatic vessel in developing heart^11–14^. In post-natal development, they seem to be required for proper mammary epithelial remodelling ^15,16^, and brain development ^17,18^. In vitro models would to experimentally study macrophages in human developmental context are currently lacking. A previous attempt at engineering human intestinal tissue with macrophages did not retrace the principles of development in its tissue derivation nor macrophage incorporation, where macrophages were injected into the lumen^22^.

A less understood aspect of macrophages is their metabolic regulation of peripheral cells. They have been shown to regulate insulin-dependent glucose metabolism of adipocytes, but also associate with cancer cells to promote their own growth^19–21^.

Here, we used previously established methods to derive human pluripotent stem cell derived intestinal organoid, which have shown to recapitulate developing intestinal architecture, along with embryonic-like macrophage^23,24^ to engineer an in vitro model

Here, we refine previous models of pluripotent stem cell derived human intestinal organoids (HIO) by incorporating macrophages and gain insight into their acquisition of antigen presentation profile as well as regulation of tissue metabolism.

## Results

### Derivation of human intestinal organoids with macrophages (HIO/Mac)

Embryonic macrophages begin to migrate and populate the organism early in the development^25,26^. We postulated macrophages would migrate and populate the intestinal organoid beyond a certain developmental time point; thus, we inquired when intestinal macrophages are first observable. In mice, AIF1 ^+^ macrophages were first observed in the mid-hindgut at E10.5, which approximately corresponds to 30 days post conception (day 30) in human development or Carnegie stage 13 (CS13)^27^ (Figure S1a-d, Figure1a). In humans, we identified macrophages in a single-cell RNA-sequencing (scRNAseq) dataset of day 47 fetal proximal intestine, the earliest dataset reported to date^28^(Figure1h-h”, Figure S1e,f). The results indicate that macrophages populate both the mouse and human intestine at an early embryonic stage. Based on these results, we estimated that the first macrophage occupation of the human intestine would be approximately at day 30.

**Figure 1.**
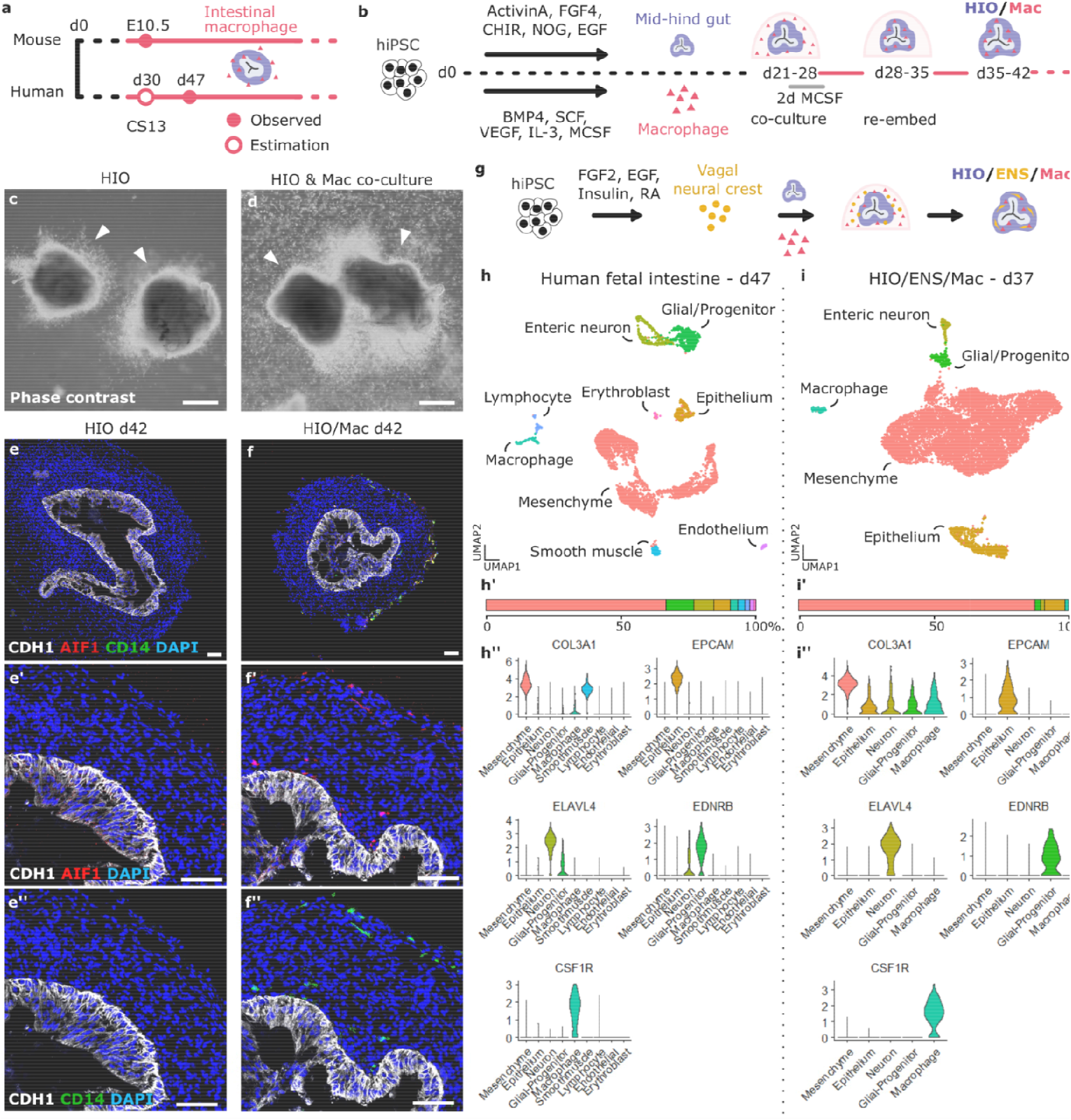
Derivation and composition of human intestinal organoid (HIO) with macrophages. **a,** Timeline of macrophage habitation of mid-hindgut in mouse and human. d, days post conception; E, embryonic day; CS, Carnegie stage. **b,** Overview of the derivation of HIO with macrophages (HIO/Mac). hiPSC, human induced pluripotent stem cell; CHIR, CHIR99021. **c,d,** Phase contrast image of day 28 HIO alone (c) or HIO in co-culture with hiPSC-derived macrophages (Mac) (d). Arrowhead, HIO. **e-f”,** Immunofluorescence of HIO (e-e”) and HIO/Mac (f-f”) for markers of epithelium (CDH1), macrophage (AIF1, CD14), and nucleus (DAPI). **g,** Schematic of the derivation of HIO with enteric nervous system (ENS) and macrophages (HIO/ENS/Mac). **h,i,** Uniform Manifold Approximation and Projection (UMAP) plot of single-cell RNA sequencing (scRNAseq) of day 47 human fetal proximal intestine (h) and day 37 HIO/ENS/Mac (i). **h’,i’,** Percent cell type composition of each dataset (h,i). **h”,i”,** Violin plot of a representative gene used to identify the cell type of each cluster in UMAP (h,i). Scale bars, 0.5mm (c,d), 75μm (e-f”).

HIOs and macrophages were derived from human induced pluripotent stem cells (hiPSC) based on previously described methods with minor modifications^23,24,29^. We used macrophage derivation which recapitulates early embryonic macrophage ontogeny^30^. Day 21-28 HIOs were used which roughly corresponds to day 30 of our previous estimation taking inner cell mass formation (~7days since conception) into account. HIOs were co-cultured with the macrophages in a three-dimensional Matrigel droplet for 7 days, during which macrophages in the periphery migrated into HIOs (Figure1c,d, Figure S1i). HIOs with macrophages (HIO/Mac) were then transferred into a new Matrigel droplet and cultured in the absence of peripheral macrophages for 7 days (Figure1b). As expected, CD14^+^/AIF1^+^ macrophages were present in HIO/Mac but not in HIO (Figure1e-f”). Furthermore, macrophages persisted within the HIO and proliferated (Figure S1e,e’). Similar to E15.5 mice, macrophages in the HIO either associated tightly with the epithelium or were found within the surrounding mesenchyme (Figure S1g,h). The procedure was successful with macrophages derived from three hiPSC-lines and HIOs from two hiPSC-lines in all instances, but with somewhat varying efficiencies (data not shown).

### Cellular composition of the intestinal organoids

We first examined the cellular composition of the organoid. Neurons of the enteric nervous system (ENS) localize closely with macrophages and together regulate peristalsis and immune response^8–10^. To increase the complexity of HIO/Mac for characterizations downstream, we derived HIO/Mac with ENS (HIO/ENS/Mac) by incorporating hiPSC-derived vagal neural crest cells, precursors to ENS^24^ (Figure1g). We then performed scRNAseq on HIO/ENS/Mac and compared it to the day 47 fetal intestine dataset. Presumptive cell identities were assigned to unsupervised clusters based on known gene markers (Figure1h”,i”). Like the fetal intestine, the organoid consisted of mesenchyme, epithelium, enteric neurons, glial-like progenitors, and macrophages. In contrast to the fetal intestine, the organoid did not yet develop any distinct smooth muscle. Expectedly, the organoid lacked endothelium, lymphocytes, and erythroblasts (Figure1h-i”). Together, the results indicate that the HIO/Mac represents an early embryonic intestine and consists of expected cell types, comparable to early fetal intestine.

### Macrophage recruitment and retention

CSF1 (MCSF) is a crucial regulator of macrophage differentiation, survival, and proliferation, which is present in circulation and produced in local tissues^31,32^. Since macrophages occupied and proliferated within the HIO/Mac, we suspected a local source of CSF1. Mesenchymal cells were the major expresser of *CSF1* in the organoid and in day 47 to 127 fetal intestines single-cell datasets. *CSF1* was also expressed in lymphocytes (T and NK), and endothelial cells in the fetal intestines, though these cells represented a much smaller proportion (Figure S2). We have previously supplemented the media with 100ng/ml of MCSF; however, these results suggested that the macrophages might not continuously need external MCSF. To the point, we found that 20ng/ml MCSF during the first two days of the co-culture was sufficient to derive HIO/Mac and the addition of MCSF was not required to maintain the organoid (Figure1b). In conclusion, we identify mesenchymal cells as the main producer of CSF1 in the developing intestine, replicated in HIO, and find that this local production is sufficient to maintain the macrophage population.

CSF1 is also a chemoattractant for macrophages. We sought to further understand signaling pathways involved in the macrophage recruitment to the organoid. To this end, we looked at a curated list of ligand-receptor pairs known to be involved in macrophage recruitment. Genes commonly upregulated in inflammatory conditions, such as *CCL2* and its receptor *CCR2*, showed little to no mutual expression in the HIO cells and macrophages, respectively. On the other hand, *CX3CR1-CX3CL1* and *CSF1-CSF1R* that are known effectors of macrophage recruitment and/or survival during the development, showed high mutual expression (Figure S3a)^33–35^. Similar results were observed in quantitative PCR (qPCR) of macrophages that were yet co-cultured with the HIO (Figure S3b). In conclusion, the recruitment of macrophages to HIOs seems to primarily involve developmentally relevant signalling rather than inflammatory pathways.

### Effect of enteric neurons in macrophage recruitment

A previous report suggested that enteric neurons are the main source of CSF1 in adult mouse intestines^10^. Conversely, our results indicated little *CSF1* expression in enteric neuron in the organoid and in day 47 to 127 fetal intestines (Figure S2). To directly test if enteric neurons affect macrophage migration and survival in the developing intestinal organoid, we derived HIO/Mac with or without the ENS. Presence of enteric neurons within the HIO did not affect the number of macrophages in the HIO (Figure S3c,d). This observation supports a previous report that the lack of enteric neurons in neonatal *Ret*^-/-^ mice and children with Hirschsprung’s disease does not affect the number of intestinal macrophages^25^. In conclusion, our results substantiate that enteric neurons do not affect the establishment of intestinal macrophages in early development.

### Macrophages migrate to the wound site upon injury

We wanted to test if macrophages in the organoid displayed behaviors of *in vivo* macrophages. A distinct feature of macrophages is their recruitment and response to tissue injury, also during early development^36,37^. To test if organoid macrophages can migrate to the injury site, we derived macrophages from eGFP-tagged hiPSC (hiPSC^eGFP^) and tracked their movements within the organoid with time-lapse imaging after a puncture injury. During the 12 hours post-injury, macrophages migrated towards the injury site, whereas the movements of uninjured organoids’ macrophages were not concerted. Assessment of their relative distance to the injury and the chemotactic precision index, which quantifies directional movement, supported this observation (Figure S4). In conclusion, we show that macrophages in the early stage *in vitro* organoid recapitulate the recruitment of macrophages to the embryonic injury observed in mice.

### Macrophages localize to intestinal micro-anatomical niches in xenograft-matured organoids

Macrophages localize to specific micro-anatomical niches within the intestine, such as villi, crypts, and enteric ganglia^5,6,8–10,38^. To test if macrophages in HIOs localize to these niches, HIO/ENS/Macs were grafted to immunodeficient *non-obese diabetic/Prkdc^SCID^/Il2rg^null^* (NSG) mice for 8-12 weeks to facilitate further development, forming more complex tissue structures^39^ (Figure2a). Furthermore, macrophages were derived from hiPSC^eGFP^ line to trace their origin. The grafted organoids maintained CDX2^+^ intestinal epithelial identity and developed crypts, villi, smooth muscle, and enteric ganglia^40^. eGFP^+^ macrophages also self-maintained over the course of engraftment and established in micro-anatomical niches of grafted organoids (Figure2b,c, Figure S5a).

**Figure 2.**
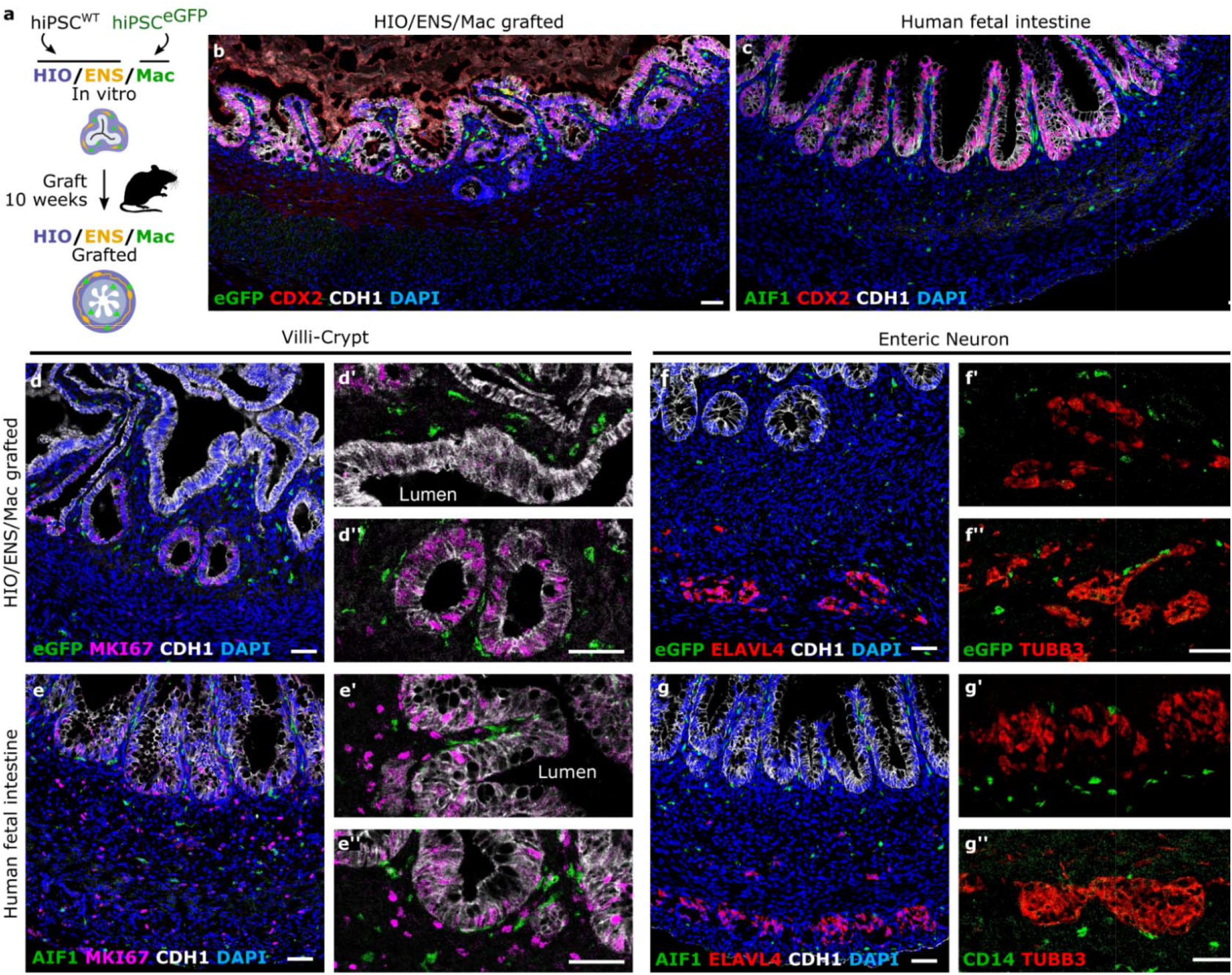
Macrophages localize to intestinal micro-anatomical niches in xenograft-matured intestinal organoid. **a,** Strategy to track macrophages by deriving it from enhanced green fluorescent protein (eGFP) tagged hiPSC (hiPSC^eGFP^) and further differentiation and growth of the organoids by xenografting to the immunodeficient (NSG) mice. **b-g”,** Representative confocal microscopy images of day 112 grafted HIO/ENS/Mac and day 119 human fetal proximal intestine. Immunofluorescence for intestinal epithelium specific identity (CDX2, CDH1) (b,c), localization of macrophages (eGFP or AIF1) within the villi (d’,e’), adjacent to intestinal crypts (MKI67, CDH1) (d, d”,e,e”), and enteric ganglia (TUBB3, ELAVL4) (f-f”,g-g”). Nuclei (DAPI). Scale bars, 50μm.

In more detail, at the epithelium, macrophages positioned flat on the MKI67^+^CDH1^+^ epithelial crypt cells and congregated within the villi, comparable to the distribution and morphology seen in day 119 human fetal intestinal macrophages and in mouse^9^ (Figure2d-e”). Intestinal monocytes/macrophages near the epithelium phagocytose lumenal content, such as bacteria^2–4^. To test if macrophages in the organoid display such transepithelial phagocytic activity, we injected Escherichia coli (E.coli) particles conjugated to pH-sensitive fluorescent dye into the lumen of grafted HIO/Mac^eGFP^. Confocal microscopy images showed that eGFP^+^ macrophages near the epithelium internalized the particles, indicating transepithelial phagocytic activity (Figure S5c). Furthermore, intestinal macrophages were shown to regulate epithelial differentiation and proliferation in adult mice^5–7^. In particular, macrophage ablation decreased the number of proliferating crypt epithelial cells^6^. To test if macrophages affect the crypt during development, we grafted HIO/ENS and HIO/ENS/Mac in parallel for comparison. Intriguingly, the number of proliferative cells in the epithelium were lower without the macrophages (Figure S5d,e). We must note here GFP^-^/AIF1^+^ macrophages were found in both HIO/ENS and HIO/ENS/Mac, indicating that macrophages from the NSG mice populated the grafted organoids (Figure S5b,b’). Previous reports indicate that macrophages of NSG mice display a delay in maturation and defects in immune function, but is unclear if their homeostatic capacities are affected ^41–45^. In this case, hiPSC-derived human macrophages seem to exert an effect that mouse macrophages cannot replicate. In conclusion, grafted organoids recapitulate macrophages localization in the intestinal mucosa and their transepithelial phagocytosis. Furthermore, hiPSC-derived macrophages seem to affect the proliferation of epithelial crypts in the developing organoid.

At the enteric ganglia, organoid macrophages localized adjacent to the ELAVL4^+^ and TUBB3^+^ neurons, resembling neuron-associated intestinal macrophages in day 119 human fetal intestine and adult mice^9,10^(Figure2f-g”). Cell ablation studies indicate that intestinal macrophages of adult mice regulate peristalsis^9,10^. To test if macrophages affect organoid contraction, we recorded the isometric force of whole grafted organoids with or without macrophages. However, we did not observe significant differences in contractility between the two conditions (Figure S5f-i). We do not draw any conclusion on peristalsis since mouse macrophages may have compensated for the lack of iPSC-derived macrophages. In short, macrophages in the organoid associated with enteric ganglia as observed in the fetal and adult mouse intestine.

### Transcriptomic comparison of the intestinal organoid’s and fetal tissue macrophages

Macrophages/monocytes that populate the intestine further differentiate to fulfill the locale-specific function^9,25,46^. We examined with scRNAseq if organoid macrophages differentiate in response to their tissue environment by comparing their transcriptional changes to that of human fetal intestine and distal lung macrophages^28,47^. Macrophages were subset from day 80 and day 127 human fetal proximal intestine and distal lung, and for day 37 *in vitro* organoid and day 121 grafted HIO/ENS/Mac^eGFP^ datasets. In detail, macrophage clusters from the unsupervised clustering of each dataset were further selected for *CSF1R* and *CD14* expressing cells (Figure3a, see Methods). Differentially expressed genes between the later and earlier time points of the intestinal and distal lung macrophages showed distinct upregulation profiles (Figure3b,c). At large, intestinal macrophages increased antigen presentation-related gene expressions (HSPs, HLAs, *TMEM176A, TMEM176B*), whereas distal lung macrophages upregulated genes that were previously known to regulate lung morphogenesis (*SCGB3A2, ADH1B, RARRES2*) and extracellular matrix protein enriched in the lung (*FBLN1*)^48–53^ (Figure3b,c,e,h).

**Figure 3.**
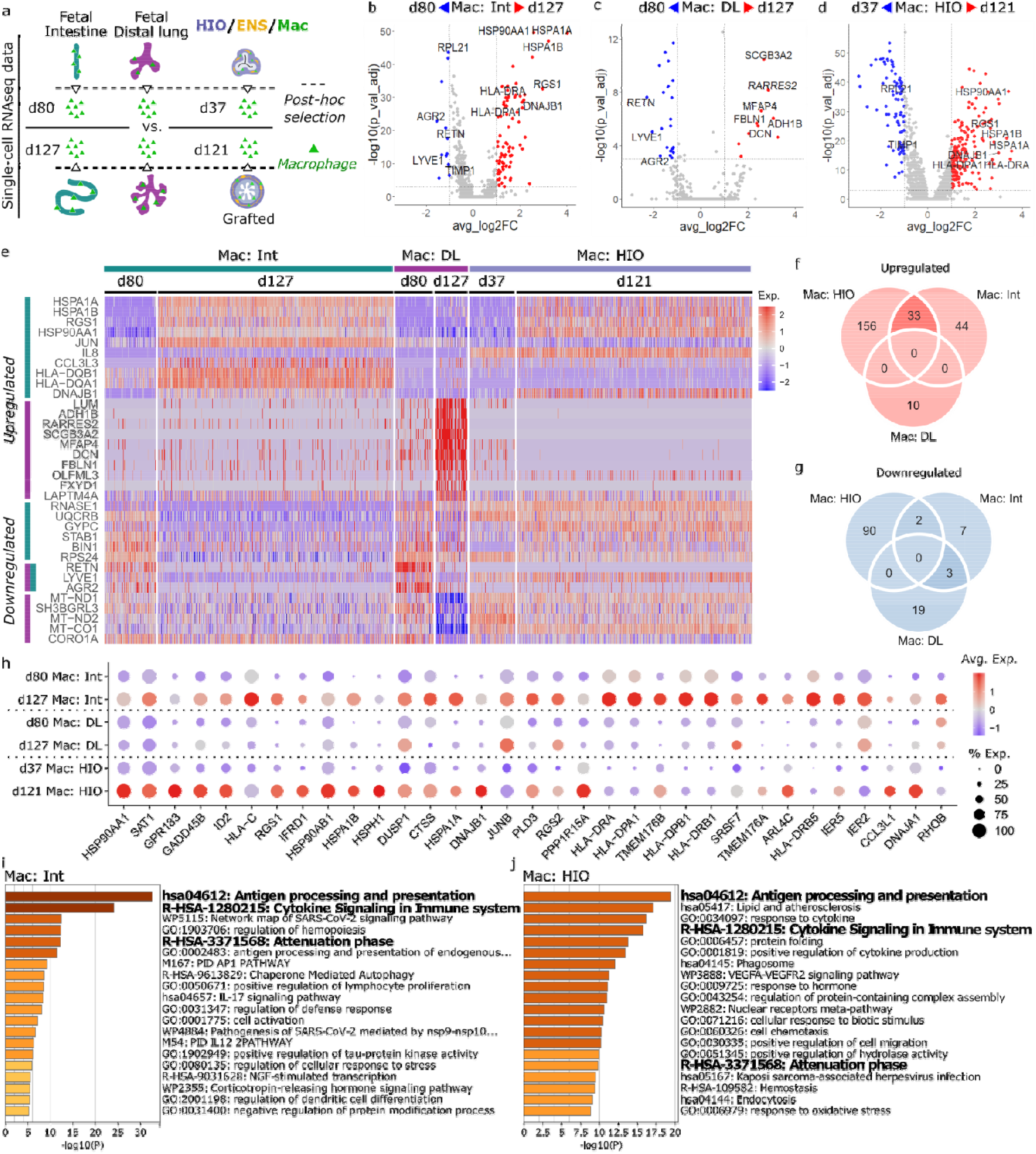
Intestinal organoid macrophages acquire transcriptomic profile alike fetal intestinal macrophages. **a,** Overview of post hoc selection of macrophages from organoid and fetal tissue scRNAseq datasets. **b-d,** Volcano plots of differentially expressed genes in the macrophages between late and early developmental time points from: fetal proximal intestines (b), distal lungs (c), and organoids (d). Threshold of discovery (dotted line), log2fold change > 1, adjusted p-value < 0.001, Wilcoxon rank sum test. Macrophage, Mac; Intestine, Int; Distal lung, DL. **e,** Heatmap of top ten upregulated and downregulated genes found in the fetal proximal intestine (b), and distal lung (c) in the order of fold change. Number of cells in ‘Mac: HIO’ were down sampled for visualization. Exp, scaled expression level. **f,g,** Venn diagram of the number of upregulated (f) and downregulated genes (g) from (b-d). **h,** Dot plot of the 33 commonly upregulated genes between intestinal and organoid macrophages (f). Avg. Exp., average expression; % Exp, percentage of cells expressing the gene. **i,j,** Gene ontology annotation generated with all upregulated genes of intestinal (i) and organoid macrophages (j).

Differentially regulated profiles were then compared to that of organoid macrophages. Heatmap of top ten differentially expressed genes from intestinal and distal lung macrophages demonstrated that organoid macrophages (Figure3d) shared many upregulated genes with intestinal but not with distal lung macrophages (Figure3e). Specifically, 33 out of 77 upregulated genes were shared between intestinal and organoid macrophages (Figure3f,h), with significant enrichment of ‘Antigen processing and presentation’ in intestinal and organoid macrophages (Figure3i,j, Figure S6b). Genes only upregulated in intestinal macrophages included additional HLA genes (Figure S6a,c). Comparisons of additional fetal intestinal datasets with varying developmental intervals showed similar signature where antigen-presentation genes are upregulated (Figure S6d,e,f). Downregulation profiles between intestinal, distal lung, and organoid macrophages (*LYVE1, RETN, AGR2, TIMP1*, and *RPL21*) did not return discernable interpretation from gene ontology (Figure3b,c,g). In conclusion, we demonstrate that macrophages in the organoid undergo similar tissue-specification as that of the fetal intestine at a transcriptional level.

### Macrophages attenuate mesenchymal cell glycolysis in developing intestinal organoids

To further investigate the roles of macrophages in early intestinal development, we performed scRNAseq on organoids with or without macrophages (two HIO/ENS pairs and one HIO pair) and compared the transcriptomic difference in different cell types (Figure4a). Differentially expressed genes common in all three sample pairs were present in the mesenchyme and epithelium (Figure4b,c). The most discernable gene ontology annotation among these results was the downregulation of glycolytic enzymes (*ENO1, LDHA, PFKP, PGK1*, and *TPI1*) in mesenchyme, suggesting a decrease in their glycolysis when macrophages are present (Figure4d,f). Macrophages were previously shown to regulate glucose uptake and metabolic response to cold in adipocytes^20,21^. They were also shown to upregulate glycolysis in cancer cells and promote tumor growth^19^. Mapping protein-protein association of the downregulated mesenchymal genes revealed a network of the aforementioned glycolytic genes as expected, but also their associations with *MIF* (Figure4e). *MIF* was previously shown to promote glucose uptake and glycolysis of muscle and cancer cells, and shown to be expressed in intestinal epithelium^54–56^. *MIF* was also downregulated in the organoid epithelium^56^ (Figure4g). In conclusion, we show that macrophages reduce glycolytic gene expression of the mesenchymal cells in the organoid.

**Figure 4.**
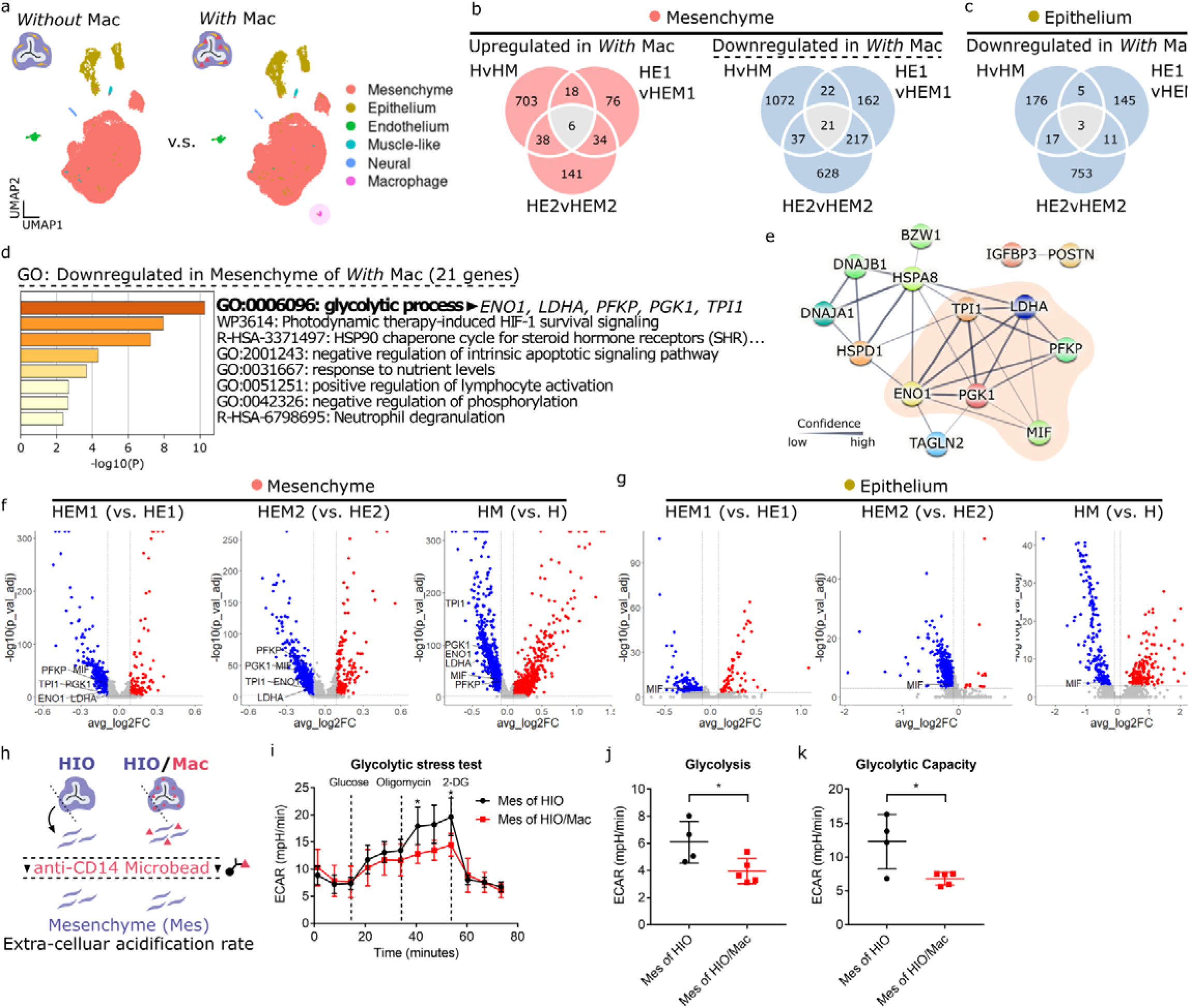
Macrophages attenuate mesenchymal cell glycolysis in developing intestinal organoids. **a,** Representative UMAP plots from scRNAseq datasets of organoids without or with macrophages (Mac). Merged dataset of HIO/ENS’s and HIO/ENS/Mac’s **b,c,** Venn diagram of the number of differentially expressed genes in organoids *With* macrophages against organoids *Without* between each sample. Commonly differentially expressed genes were found in mesenchyme (b) and epithelium (c). HvHM, HIO vs. HIO/Mac; HE1(2)vHEM1(2), HIO/ENS1(2) vs. HIO/ENS/Mac1(2). Each sample is a pool of 4-5 dissociated organoids. Threshold of discovery: adjusted p-value < 0.001, Log2fold change > 0.09, Wilcoxon rank sum test. **d,** Gene ontology analysis on the 21 genes downregulated in mesenchymal cells of the organoid due to macrophages (b). **e,** Protein-Protein association map of the 21 genes (b) from STRING. Only the genes with at least one association is displayed. **f,g,** Volcano plots annotated with genes involved in glycolytic process as indicated by (d) and protein-protein association map (e). Dotted lines: adjusted p-value < 0.001, Log2fold change > 0.09 & < −0.09, Wilcoxon rank sum test. **h,** Schematic of mesenchymal cell preparation for glycolytic stress test from *in vitro* organoids. **i,** Extracellular acidification rate (ECAR) measurement in glycolytic stress test. Oligomycin (ATP synthase inhibitor). 2-DG, 2-Deoxy-D-glucose (competitive inhibitor of glucose). **j,k,** Quantification of glycolysis (j) and glycolytic capacity (k) as indicated by the glycolytic stress test (i). See Methods for calculation. Result of one experiment (i-k). Mean & s.d (i-k). p < 0.0001, Two-way ANOVA; p = 0.0311, p = 0.0289, post-hoc: Sidek-Bonferroni (i). Each data point represents a technical replicate well. p = 0.0360 (h), p = 0.0199 (i), student’s t-test.

We used metabolic-flux assay (Seahorse XF) to test whether macrophages downregulate glycolysis in HIO as predicted by scRNAseq. Mesenchymal cells were dissected and isolated into single cell suspension from HIO or HIO/Mac. Macrophages were removed from the cell suspension with anti-CD14-mediated magnetic separation (Figure S7a-e). Extracellular acidification rate (ECAR) was measured to gauge the glycolytic activity of mesenchymal cells (Figure4h). In the glucose-depleted state, both conditions showed comparable ECAR (Figure S7f). Introduction of glucose induced an increase in ECAR as expected; however, mesenchymal cells from HIO/Mac showed a smaller increase indicating a lower level of glycolysis (Figure4i,j). Subsequent inhibition of mitochondrial respiration with oligomycin, necessitating the cells to use glycolysis-driven ATP generation, showed that the glycolytic capacity was also lower in mesenchymal cells from HIO/Mac (Figure4i,k). Lastly, inhibition of glucose-uptake by 2-Deoxy-D-glucose (2-DG) decreased the ECAR again to a comparable level (Figure4i). Thus, metabolic flux assay supports our transcriptomic result that macrophages reduce glycolysis of mesenchymal cells.

### Macrophages regulate intestinal organoid growth

Intriguingly, organoids engrafted for 10 weeks were smaller when macrophages were incorporated (Figure5a,b,d). Organoids had been randomized for the co-culture with macrophages during their derivation and their size shortly after the macrophage co-culture was still comparable (Figure S8a). Histology did not show any difference in tissue morphology between the two conditions. In particular, we did not observe any signs of abnormal fibrosis, which could arise due to prolonged inflammatory state by pro-inflammatory macrophages and stunt growth (Figure5a’-a’”,b’-b’”). Additional grafted organoids with macrophages collected at 8 weeks and 12 weeks after engraftment were also smaller in size (Figure5c,e, Figure S8b). Exponential fit of the organoid size as function of time indicated that the rate of growth was reduced when incorporated with macrophages (Figure5f).

**Figure 5.**
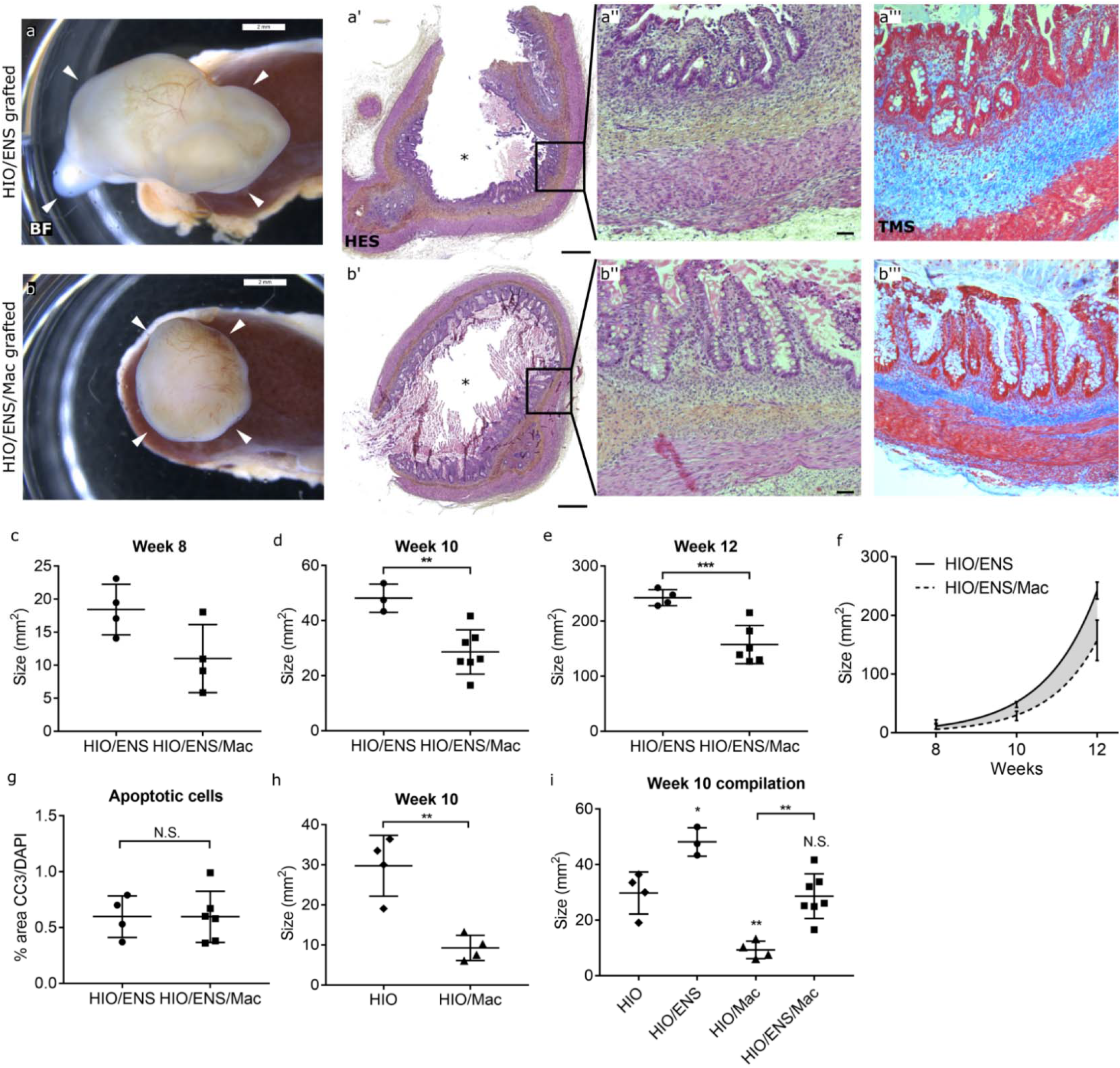
Macrophages reduce the growth of intestinal organoid without affecting tissue architecture. **a-b’”,** Representative bright field (BF) images of week 10 grafted organoids still attached to the mouse kidney at the site of the engraftment, White arrowheads point to the organoid (a,b). Hematoxylin-eosin-saffron (HES) stained sections of the grafted organoids (a’,a”,b’,b”). Trichrome Masson (TMS) stained sections (a’”,b’”). Asterisks, lumen. **c,d,e,h,** Size measured by the area of the grafted organoids from bright field images, isolated at 8 weeks (c), 10 weeks (d,h), and 12 weeks (e) after the engraftment. **f,** Exponential regression of the size of the grafted organoids over time from (c-e). **g,** Level of apoptosis in grafted organoids. Quantified by calculating the area of cleaved caspase 3 (CC3) divided by the area of nuclei (DAPI) in confocal microscopy images of immunofluorescence. **i,** Sizes of all 10 week grafted organoids combined with or without ENS and/or Mac. Each data point represents an organoid. HIO/ENS, n = 4, HIO/ENS/Mac, n = 4 (c), HIO/ENS, n = 3, HIO/ENS/Mac, n = 7 (d), HIO/ENS, n = 4, HIO/ENS/Mac, n = 6 (e), HIO/ENS, n = 4, HIO/ENS/Mac, n = 6 (g), HIO, n = 4, HIO/Mac, n = 4 (h), mean & s.d, p = 0.0633 (c), p = 0.0034 (d), p = 0.0010 (e), p = 0.995 (g), p = 0.0075 (h), Welch’s t-test. p < 0.0001, two way ANOVA (f). p < 0.0001, one way ANOVA; HIOvs.HIO/ENS, p = 0.0148, HIOvs.HIO/Mac, p = 0.0038, HIOvs.HIO/ENS/Mac, p = 0.9930, HIO/ENSvs.HIO/Mac, p < 0.0001, HIO/ENSvs.HIO/ENS/Mac, p = 0.0045, HIO/Macvs.HIO/ENS/Mac, p = 0.002, postdoc: Tuckey (i). Scale bar, 2mm (a,b), 0.4mm (a’,b’), 50μm (a”,a’”,b”,b’”).

Macrophages are professional phagocytes; thus, we next tested if the smaller size in grafted organoids with macrophages is due to the removal of apoptotic cells. Level of immunofluorescence signal for apoptotic cells; however, were comparable (Figure5g). Thus, the size difference is not due to the removal or accumulation of apoptotic cells.

The report that described the HIO/ENS derivation noted that organoids combined with ENS had higher EGF expression via bulk-RNA sequencing, though size difference was not noted^24^. Furthermore, macrophages in the brain (microglia) were shown to phagocytose neuronal processes and required for proper brain development and morphology^17,18^. Intestinal macrophages may suppress overexpansion of enteric neurons and in turn inhibit tissue overgrowth. Thus, to test a part this concept, we next hypothesized that the reduced growth of organoids by macrophages is dependent on the presence of ENS. Our experiments showed that ENS did have positive growth effect on grafted HIO; however, it also revealed that macrophages limited the organoid growth independent of ENS (Figure5h,i, Figure S8c). We thus conclude that ENS and macrophages positively and negatively regulate organoid growth, respectively, and that neither of the effects are necessarily dependent on each other. The independent effect of macrophages on organoid growth may be due to the reduction of glycolysis observed here; however, further investigation is required.

## Discussion

We demonstrate a human intestinal tissue model with macrophages. The macrophages localized to known intestinal micro-anatomical niches in xenograft-matured organoids. They also acquired a similar transcriptional profile to fetal intestinal macrophages. Functionally, they phagocytose lumenal antigens and migrate to wounds upon injury. Furthermore, experimental studies with the model suggest that macrophages regulate the metabolism of the developing intestine and organ growth.

Modularity of the HIO/Mac, as demonstrated by vagal neural crest cell or macrophage exclusion experiments here, is particularly useful when cellular composition is an experimental variable. Similar cell exclusion experiment should be useful to examine specific effects of embryonic macrophages in tissue development, such as their role on vascular and lymphatic development^10,13,14^.

Macrophages are required for proper regeneration in adult^36,57^. Scarless wound healing in the fetus can be an intriguing avenue to study mammalian wound healing. Further adaptation of the method described here should be useful to explore the mechanisms and limitations of wound healing and how macrophages react in fetal injury or infection.

The organoids were kept in aseptic conditions during both *in vitro* culture and engraftment. Thus, the induction of the antigen-presentation profile in organoid macrophages, also observed in fetal intestine, is occurring in the absence of microbiota. This suggests that intestinal tissue environment itself contributes to this specification, and is already present in prenatal development.

Unexpectedly, we find that HIO with macrophages are smaller than HIO without. Build-up of apoptotic cells due to lack of macrophages does not seem to be the cause of the size difference. Our scRNA-seq and metabolic analysis show that this most likely parallels a decrease in glycolysis and glycolytic capacity, which then in turn limits organ size. This effect occurs independent of the positive regulation of organ growth by the ENS. Positive trophic effect of the ENS has also been observed recently in gastric organoids. In summary, our results suggest that these two distinct cell populations function through non-overlapping mechanisms which can affect tissue/organ growth during embryonic development.

In conclusion, we report an improvement to an existing intestinal tissue model, which now harbor macrophages during key developmental stages. We expect the model to facilitate the study of macrophage specifications and functions in human tissues and diseases.

## Acknowledgement

We also recognize and thank the following individuals and organizations. CHUSJ core facilities: iPSC core facility for reprograming of cells and gene editing, Basma Benabdallah; Cytometry, Ines Boufaied; Imaging by microscopy, Elke Kuster. Sabine Herblot from Michel Duval lab provided the human dendritic cell. Sylvain Chemtob lab offered the use of isometric force measurement apparatus

## Author contribution

A.S. designed the study. R.S. with A.S. and S.L. derived the macrophages. Y.L. performed surgical procedures in mice. S.L. and P.V. prepared the libraries for scRNAseq. A.S. with H.A. performed bioinformatics analyses. T.A. performed the glycolytic stress test. M.C. and A.S. performed image analyses. J.G. and N.P. collected the fetal tissues. A.S. conducted the other tasks. A.S. wrote the paper with contributions from the authors. G.A. supervised all aspects of the study and provided funding.

## STAR Methods

### Cell lines and tissues

The use of human pluripotent stem cells was approved by the ethical committee at the participating institution CHU-Sainte Justine, Montreal QC, Canada (CER#2287). The hiPSCs and their derivations were routinely tested for mycoplasma (LT07-118, Lonza) and tested negative.

Human fetal (17-21 weeks) tissue was obtained after written informed consent and was approved by the ethical committee of CHU Sainte-Justine, Montreal QC, Canada (CER#2126).

The animal procedures were approved by the animal committee of CHU Sainte-Justine, Montreal QC, Canada (CIBPAR#2021-3202, 2022-3784).

### Pluripotent stem cell culture

Human induced pluripotent stem cells (hiPSC) were generated at CR-CHUSJ Stem Cell core or in-house. The hiPSC lines SJi3252C2 and SJ3013C2 were derived from human fibroblasts using Cytotune 1.0^™^ (Invitrogen) and were previously characterized^58^. EU03.C2, and EU148.C5 were derived from human PBMC with Cytotune™ 2.0 (A16517, Invitrogen). The hiPSC^eGFP^ line (SEC61B-GFP hiPSC, AICS-0010) was acquired from Allen Institute for Cell Science^59^. The hiPSCs were cultured in hypoxic condition (5%CO_2_, 5%O_2_, 37°C incubator) until passage 15-20, otherwise all cells and derivatives were cultured in normoxic condition (5%CO_2_, 37°C incubator). They were cultured with mTeSR1 (85850, StemCell Technologies) with 1X penicillin-streptomycin (450-201-EL, MultiCell) and hESC-qualifed Matrigel (354277, Corning). Matrigel was coated onto Nunc Delta surface plates (14-832-11, Thermo Scientific) as per manufacturer recommendation. The cells were passaged as small clusters using 0.5mM ethylenediaminetetraacetic acid (EDTA) in phosphate buffered saline (PBS). The cells were cryopreserved with NutriFreez™D10 (05-713-1E, Biological Industries) as per manufacturer recommendation.

### Human intestinal organoid derivation

Human intestinal organoids (HIO) were derived with the hiPSC-lines SJi3252C2 or SJ3013C2 as previously described with minor modifications^24,60,61^. Briefly, 85% confluent hiPSCs in a 6 well plate were passaged with EDTA and seeded 1/14-1/16 of a cell suspension per single well of 24 well plate. The cells were fed mTeSR1 daily for two days or until the confluency reached 80%. On the first day, the media was changed with endoderm differentiation media base (EDM-base): RPMI1640 (11875-093, Gibco), 1X pen-strep (15015067, Wisent), 1X non essential amino acid (11140050, Gibco) with 100ng/ml Activin A (338AC010, R&D systems). On the second day, the cells were fed EDM-base supplemented with 100ng/ml Activin A and 0.2% FBS (HyClone™, Fisher Scientific). On the third day, the cells were fed the same as the second but with 2% FBS. At the end of the third day, the monolayer expressed definitive endoderm markers SOX17 and FOXA2 (Figure S9a). The fourth day, the confluent monolayer of cells were fed with mid-hindgut differentiation media (MHDM): EDM-base, 2% FBS, 500ng/ml FGF4 (235F4025CF, R&D systems), 3μM CHIR99021 (S1263, Selleckchem). The cells were fed MHDM daily for a total of 4 days. At the end of mid-hindgut differentiation, the free-floating spheroids were collected and suspended in Matrigel (354234, Corning) supplemented with 1X B27^™^ supplement (17504044, Gibco) and 100ng/ml EGF (236EG200, R&D systems; AF-100-15, PeproTech). The Matrigel suspension were plated as a droplet with 15-20 spheroids per droplet on a plate (14-832-11 or 130184, Thermo Scientific) and polymerized at 37°C for 10 min. Note that some tissue culture plate types are not suitable for Matrigel droplet formation. The spheroids were fed intestinal basal media (IBM): Advanced DMEM-F12 (12634-010, Gibco), 1X B27 (17504044, Gibco), 1X Glutamax (35050061, Gibco), 1X pen-strep, 15mM HEPES (15630080, Gibco) supplemented with 100ng/ml EGF and 100ng/ml Noggin (6057NG025CF, R&D systems) for four days changing the medium every 48 hours. They were then fed IBM supplemented with 100ng/ml EGF (IBMe) every 48 hours. Two weeks after the spheroid collection, the organoids were passaged by manually separating from each other with sterile syringe needle and re-embedding in the Matrigel droplet as before. From this point onward, the organoids were fed IBMe and passaged every two weeks until the experiment. The organoids were positive for intestine-specific epithelial marker CDX2 (Figure S9b).

### Vagal neural crest derivation

Vagal neural crest cells (VNCC) were derived from the hiPSC-line SJ3013C2 as previously described with minor modifications^24,62^. Briefly, In a 6 well plate, small colonies of hiPSC were seeded at low density (1/80 of a confluent well of 6 well plate) and grown for 5 days. On the first day of differentiation, the colonies were lifted with 500U/ml collagenase IV (17104019, Gibco) in mTeSR1 for up to 1hour in the incubator or until the colonies detached completely with gentle taps to the plate. The colonies were washed with 2ml of DMEM/F12 (319-075-CL, Wisent) three times. They were then suspended in neural induction media (NIM): 1:1 ratio of DMEM-F12 and Neurobasal media (21103049, Gibco), 0.5X B27, 0.5X N2, 5ug/ml insulin (I2643, Sigma), 20ng/ml basic-FGF (100-18B, PeproTech), 20ng/ml EGF, and 1X pen-strep with 5uM of Rock inhibitor Y27632 (S1049, Selleckchem) and transferred to non-tissue culture treated plate (08-772-51, Fisher scientific). The NIM was changed daily with decreasing Y27632 (Y27) concentration (Day 2: 2.5uM Y27, Day 3 and beyond: no Y27), for additional 5 to 6 days until the neurospheres had clear round borders. The neurospheres were then fed NIM with 2uM all-trans-retinoic acid (R2625, Sigma Aldrich) daily for two days. The neurospheres were transferred on to the fibronectin (PHE0023, Gibco) coated plate in NIM (w/o retinoic acid) and left undisturbed for 48hours. Fibronectin coated plates were prepared by incubating plastic tissue culture plates with 15ug/ml fibronectin in PBS without calcium or magnesium at 37°C overnight. Afterwards, the NIM was changed daily until neural crest cells migrated and spread out onto the plate (6-10days). Neurospheres were mechanically removed and the migrated neural crest cells were lifted as single cells with a 5 minute incubation at 37°C with 1X TrypLE (A1217701, Gibco). Cells were washed by diluting with 9ml of room temperature DMEM/F12 and centrifuging at 300G for 4min. The supernatant was removed and cells were resuspended in DMEM/F12 for co-culture with HIO or resuspended in NIM and plated back onto fibronectin coated plate and maintained until the experiment. The cells were positive for known vagal fate neural crest cell markers and gene expression (Figure S9c-e).

### Macrophage derivation

Macrophages were derived from EU03.C2 or EU148.C5 or AICS-0010 as previously described with modifications^29^. Briefly, embryoid bodies (EB) were formed as follows. The hiPSC were cultured to 85-90% confluency in a 6 well plate with mTeSR1 and Geltrex matrix (A1413301, Gibco) and divided into smaller segments by scratching the bottom of the well into around 100 squares with a 100ul tip. The segments were detached mechanically with a scraper and the cells clumps were transferred into 6 well ultra-low attachment plate (3471, Corning) with EB media: mTeSR1 supplemented with 50 ng/ml BMP4 (120-05ET, PeproTech), 50 ng/ml of VEGF (100-20A, PeproTech), 20ng/ml of SCF (130-093-991, Miltenyi Biotec) and 10uM Y-27632 (72304, Stem Cell Technologies) and cultured in a 37°C, 5%CO_2_ incubator. Every two days, half of the EB media was replaced with fresh EB media without Y-27632 for a total of 7 days. The EBs were then transferred to a 6 well tissue plate (10-15 EBs/well) in Factory EB (f-EB) media consisting of X-VIVO15 (BE04-418F, Lonza) supplemented with 100 ng/ml of M-CSF (216-GMP-500, R&D), 25 ng/ml of IL-3 (PHC0031, Gibco), 1X Glutamax, 1X pen-strep and 0.055 mM b-mercaptoethanol (31350-010, Invitrogen). The f-EB media were replaced weekly. Macrophage precursors started to emerge in the supernatant after approximately 15-20 days, reaching the maturity after 30 to 45 days in f-EBs cultures, assessed by the cell surface markers expression (CD34^neg or low^, CD14^high^, CD45^high^) through flow cytometry (Figure S9f,g). The precursors were harvested every two weeks for experimentation. Harvested precursors were cultured in RPMI media (SH3009601, Hyclone) supplemented with 100ng/ml of M-CSF, 2mM L-glutamine, 10% FBS (080150, Wisent) and 1X pen-strep for 7 days, replacing the media every two days. Differentiated macrophages were assessed with flow cytometry for the expression of macrophage markers (CD14^high^, CD206^high^ CD163^high^, CD16^high^ CD11b^high^, HLADR^neg^) (Figure S9h,i,j). Additionally, dendritic cell marker FLT3 expression in the macrophages were low to none (Figure S3a,b).

### HIO combination with VNCC and macrophage

Prior to the procedure, two to three week HIOs (since spheroid collection), differentiated macrophages, and vagal neural crest cells were derived as required. Matrigel was supplemented with 1X B27 and 100ng/ml EGF. In a 5ml tube, 50K VNCC and/or 100K macrophages were added to maximum x15 HIOs. Optimal number of VNCC and macrophages may vary depending on the cell line. The mix was briefly titurated using a 1000ul pipette-tip, kept at −20°C, where 1mm of the end of the tip was cut. The mix was then centrifuged at 300G for 3min. The HIOs and the cells were then gently resuspended and centrifuged again. Supernatant was removed as much as possible and 50ul of cold Matrigel was added. The tube was kept on a cold tube from this point on. Using a cut-pipette tip that is cold, HIO and cells were gently resuspended and the total volume was gently pipetted as a droplet on a pre-warmed 6well plate. Up to four droplets were plated per well. The plate was gently transferred to the incubator and allowed to polymerize for 10min. The well was then filled with 2ml of IBMe supplemented with 20ng/ml MCSF (Thermofisher, PHC9501; PeproTech, 300-25). After 48hr, the medium was replaced with fresh IBMe. The medium was refreshed every 48hr. One week after the co-culture, the organoids were removed from the Matrigel with gentle tituration and dissection with sterile syringe needles. They were then re-embedded in new Matrigel droplets. HIOs were maintained with IBMe on the same feeding interval as before for another week before experimentation unless specified otherwise.

### Human dendritic cell

Human dendritic cells used as positive control for FLT3 qPCR were derived from purified cord blood CD34+ progenitors as previously described^63^.

### Immunocytochemistry

Cells were grown on plastic coverslips (174969, Thermo Fisher). At room temperature, cells were washed once with PBS and fixed in 4% paraformaldehyde (PFA) for 10min. PFA was removed and washed three times with cold PBS 5 minutes each. Cells were then incubated in blocking buffer (0.5% Triton X-100 and 10% donkey serum in PBS) for 1hr. Blocking buffer was removed and cells were incubated with primary antibody in blocking buffer at 4°C overnight. The next day, primary antibody buffer was removed and cells were washed in cold PBS four times 4minutes each then incubated in secondary antibody in blocking buffer for 1hour at room temperature in the dark. The secondary antibody buffer was removed and cells were washed in PBS four times 4minutes each. Cells were then stained with DAPI (D-21490, Invitrogen) and washed once with distilled water. The coverslip was mounted on a 1.5 coverslips with aqueous mounting medium (S3023, Dako) as per manufacturer recommendation before imaging.

### Immunofluorescence and histology

*In vitro* organoids, E9.5, and E10.5 mouse embryos were fixed with 4% PFA for 3hours at 4°C. Fetal proximal intestine, grafted organoids (cut in half), E12.5, and E15.5 mouse embryos (heads removed) were fixed for 12-16 hours at 4°C. The samples were washed with PBS on ice for 1hour three times with rocking and incubated in 30% sucrose in PBS at 4°C until it sank. It was then incubated with optimal cutting temperature (OCT, FSC22, Surgipath) compound at 4°C for 30 min, frozen-embedded in fresh OCT, and stored at −80°C. The samples were cut 5-10μm thickness and mounted on a charged glass slide (12-550-15, Fisherbrand). The sections were dried at room temperature for minimum 30minutes before the staining procedure. The sections were washed with PBS for 5 minutes two times and incubated in 0.5% Triton X-100 in PBS for 20 minutes. They were then incubated in the blocking buffer (10% Donkey serum and 0.3% Triton-X100 in PBS) for 30 minutes and incubated with primary antibody overnight at 4°C in a humid chamber. The next day, the slides were washed three times for 5 minutes each in PBS and incubated with the secondary antibody buffer for 1hour at room temperature. The slides were washed four times for 5 minutes each in PBS. The sections were counterstained with DAPI (D-21490, Invitrogen) for 10 minutes and washed once with distilled water. The slides were mounted with aqueous mounting medium (S3023, DAKO) with 1.5 glass coverslip and cured overnight at room temperature. The samples were then imaged with widefield (DMI8, Leica) or confocal (TCS SP8, Leica) microscope. Colorimetric staining (H&E+S and TMS) were performed by CHU-Sainte Justine pathology laboratory.

### Quantitative PCR

RNA was isolated with a DNase treatment (74104, 79254, Qiagen; Z6111, Promega) and cDNA was synthesized (18090050, Invitrogen) with oligo-dT (18418012, Invitrogen). SyberGreen (34226600, Roche) was used to quantify gene expression with Roche LightCycler 96 or LightCycler 480. A total of 15ng of cDNA and 0.3uM of each primer pair was used for a 15ul reaction as per manufacturer instruction. The primer sequences were designed using NCBI Primer-BLAST. Refer to table1 for the list of primer sequences.

### Grafting of organoids to immunodeficient mice

Two to three weeks after the macrophage co-culture, organoids with single epithelial structure were grafted under the kidney capsules of immunodeficient mice as previously described, but without the collagen encapsulation of the organoid prior to grafting^39^. The organoids were collected at corresponding time points.

### E.Coli particles injection into the grafted organoid lumen

Kidneys with the grafted HIO/Macs were surgically exposed again 10 weeks after the engraftment. With an insulin syringe, up to 50-200ul of fluid was removed from the lumen of the organoid and an equal volume of 4mg/ml of pH-sensitive E.Coli Bioparticle (P35361, Thermofisher) reconstituted in PBS was injected into the lumen. The grafted kidney was placed back in the mice with the same procedure for grafting the organoids. The organoids were collected 24 hours later, sectioned in half with a surgical scalpel, and fixed in 4%PFA for 3 hours. The tissue was processed and examined with the method as described in ‘Immunofluorescence’ but stained only with DAPI.

### Live imaging of injured HIO/Mac

HIO/Mac combined with hiPSC^eGFP^-derived macrophages were cultured in suspension in ultra low attachment plate (3471, Corning) with phenol-free IBMe for two days prior to injury. Advanced DMEM-F12 was replaced with high-glucose phenol-free DMEMF12 (D1145, SigmaAldrich) to make phenol-free IBMe. To injure the organoid, 20G blunt tip needle attached to a 1ml syringe was used to puncture the organoid at the center. The plunger was drawn slowly as pulling back to remove the punctured material. The injured organoid was then placed in a glass-bottom dish (0030 740.017, Eppendorf) with phenol-free IBMe and imaged at a 10min interval for 12-18hours with up to a 100um z-depth (TCS SP8 confocal or DMI8 widefield, Leica). The stage-top incubator was set at 5% CO2 at 37°C (iNU GSI2, Tokaihit). From the maximal projection image, cells were tracked with ImageJ plugin: Manual Tracking. The region of the injury was determined in situ from the images and the center of that region was used for downstream analysis. For controls, an arbitrary point at the center of the image was set, blinded from tracking of the cells. Directness and chemotactic precision index (CPI) were calculated as previously described^64^.

### Isometric Force Measurement

At week 12 of engraftment, HIO/ENS and HIO/ENS/Mac were isolated from the mice into Krebs buffer (NaCl 117mM, KCl 4.7mM, MgCl hexahydrate 1.2mM, NaH2PO4 1.2mM, NaHCO3 25mM, CaCl2 dihydrate 2.5mM, Glucose 11mM). Each whole organoid was then hooked at the opposite end with silk strings and transferred to organ bath chambers with 37°C Krebs buffer fed with 95% O_2_ and 5%CO_2_. The samples were then connected to the isometric force transducer and equilibrated at 1g of tension for an hour before the acquisition (Biopac MP150; Acqknowledge, Biopac). Spontaneous contractions were measured for 20-40min. The last 20min segment of regular contractions were taken for analyses. Maximal force is the highest peak. Contraction and relaxation time are averages of measure from trough to peak and peak to trough, respectively. Frequency of contraction is the number of peaks.

### Single cell RNA sequencing and analysis

For the preparation of single cell suspension, all samples were cut into <1mm pieces using a scalpel and dissociated using 1000U/ml collagenase IV (17104019, Thermofisher) in 2X TrypLE (A1217701, Thermofisher) with 10mM Glucose for 100-120minutes at 37°C. The suspension was gently titurated every 10 minutes to facilitate the dissociation. Cells were then passed through a 70um filter and ten-fold diluted in cold 1%FBS in PBS. The cells were then centrifuged 500G 5min at 4°C. The pellet was resuspended in 1%FBS in PBS and viable cell numbers were quantified with trypan blue and hemocytometer. For grafted day 121 HIO/ENS/Mac, GFP^+^ macrophages were enriched using fluorescence-activated cell sorting (FACS). The cDNA and the libraries were then generated using Chromium Next GEM v3.1 according to the manufacturer guideline (1000268, 10X Genomics; manual: CG000315 RevB) and sequenced at Génome Québec with Novaseq 6000 S4 (20027466, Illumina). Human fetal scRNAseq FASTQs were acquired from ArrayExpress: E-MTAB-8221, E-MTAB-9720^28,47^. For the comparison of human fetal and organoid datasets, both were processed with SoupX to negate the effect of ambient RNA, largely arising from the red blood cells^65^. Otherwise, FASTQs were processed with CellRanger and downstream analyses performed with R using Seurat and visualized with ggplot2^66,67^. When the sex of the samples could not be matched in differential gene expression, chromosome X and Y genes were omitted from further analyses and visualization. Gene ontology analyses were performed with Metascape and protein association analyses with STRING^68,69^.

### Glycolytic Stress Test

Mesenchyme of HIOs co-cultured with or without macrophages for 7days were dissected and dissociated into single cells with the same method as described in ‘Single cell RNA sequencing and analysis’. In order to remove the macrophages from the cell suspension, all samples were processed with magnetic cell sorting with CD14 Microbead (130-050-200, Miltenyi biotec), LS column (130-042-401), and MidiMACS Seperator (130-042-302) according to the manufacturer recommendation. Sorted cell suspension were centrifuged 400G for 5min at 4°C. Cells were resuspended in 4°C IBM, counted using hemocytometer, and 50K cells per well in 180ul volume were distributed to the Seahorse 96well microplate (101085-004, Agilent) pre-coated with hESC-qualifed Matrigel (354277, Corning) as per manufacturer instructions. The plate was then centrifuged 300G for 1min. The mesenchymal cells were incubated at 5%CO_2_, 37°C incubator for 1 hour for cell attachment. Subsequently, the culture media was replaced with 37°C assay media (DMEM 5030 media, 2mM glutamine) and equilibrated at 37°C non-CO_2_ incubator for 1 hour prior to extracellular acidification rate (ECAR) measurements with the Seahorse XFe96 Flux Analyzer. Following reagents were prepared according to the manufacturer instruction to perform the glycolytic stress test: 10mM Glucose, 5μM Oligomycin (O4876, Sigma-Aldrich), 100mM 2-Deoxy-D-glucose (2-DG, sc-202010A, Santa Cruz). Glycolytic parameters were calculated as follows: Glycolysis (maximum rate measurement before oligomycin measurement – last rate measurement before glucose injection), glycolytic capacity (maximum rate measurement after oligomycin measurement - last rate measurement before glucose injection), and non-glycolytic acidification (last rate measurement prior to glucose injection).

### Image analyses and quantification

Images were processed and analyzed using ImageJ and Cell Profiler for quantification.

## Supplemental Information

**Figure S1.**
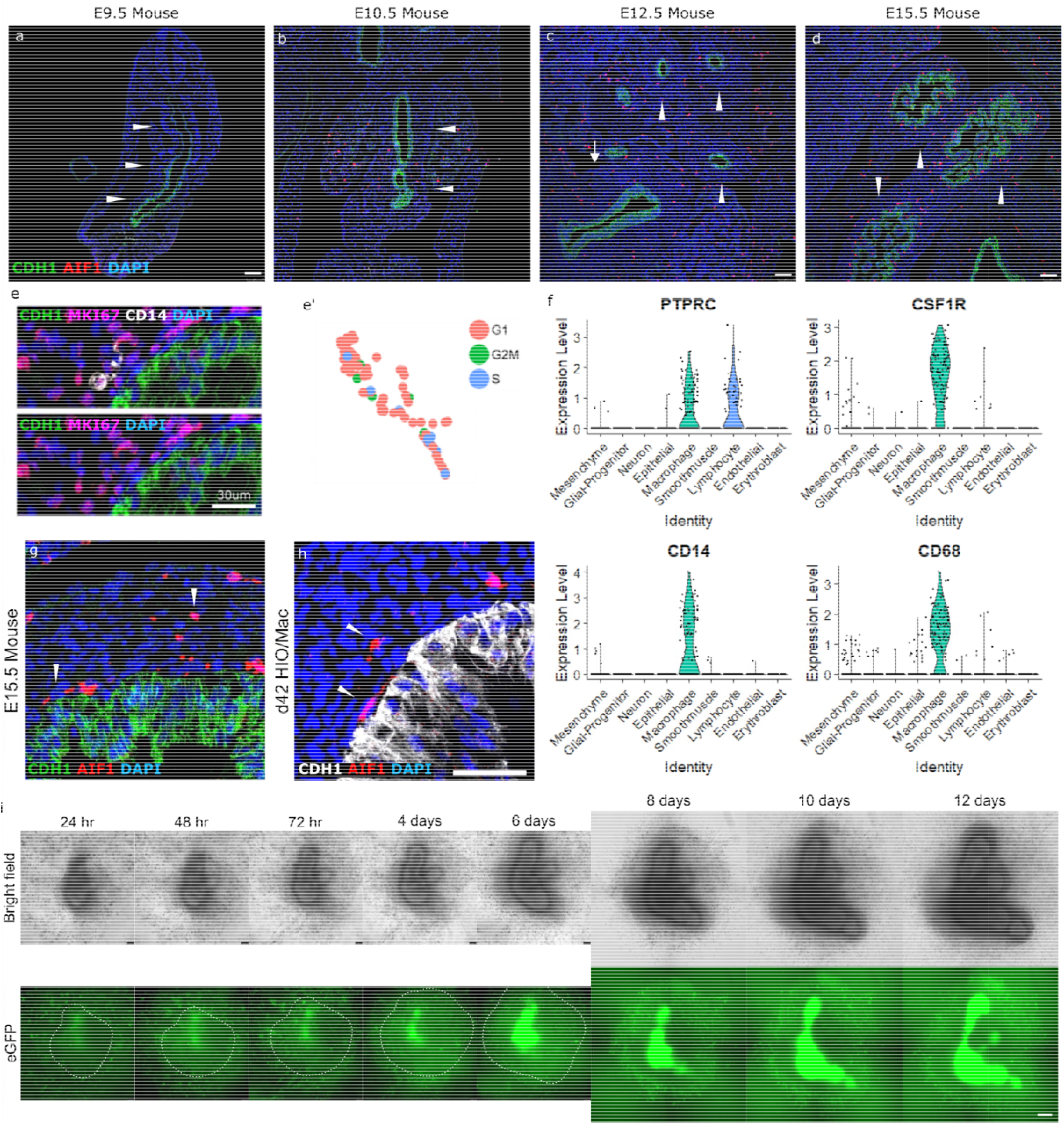
Macrophages in early embryonic mouse intestine, fetal intestine, and human intestinal organoid (HIO).**a-d,** Representative immunofluorescence images of mouse embryonic intestine for macrophages (AIF1), epithelium (CDH1), and nuclei (DAPI) at four time points of development. Arrowhead, mid-hindgut. Arrow, Foregut. **e,e’,** Representative immunofluorescence image of day 42 HIO/Mac for epithelium (CDH1), proliferation (MKI67), macrophages (CD14), and nuclei (DAPI) at four stages of development. **f,** Violin plot of gene markers used to identify the unsupervised cluster ‘Macrophage’ as macrophage. **g,h,** Representative immunofluorescence images for macrophages (AIF1), epithelium (CDH1), and nuclei (DAPI) in E15.5 mouse intestine and day 42 HIO/Mac. Arrowhead, macrophages localized at adjacent to epithelium and within mesenchyme. **i,** Bright field and GFP fluorescence images of human intestinal organoid (HIO) and macrophage co-culture over the course of 12 days. Macrophages were derived from hiPSC^eGFP^. E9.5, n=4, E10.5, n=6, E12.5, n=6, E15.5, n=6, (a-d). Scale bar, 75μm (a-d), 30μm (e), 50μm (g,h), 150μm (i).

**Figure S2.**
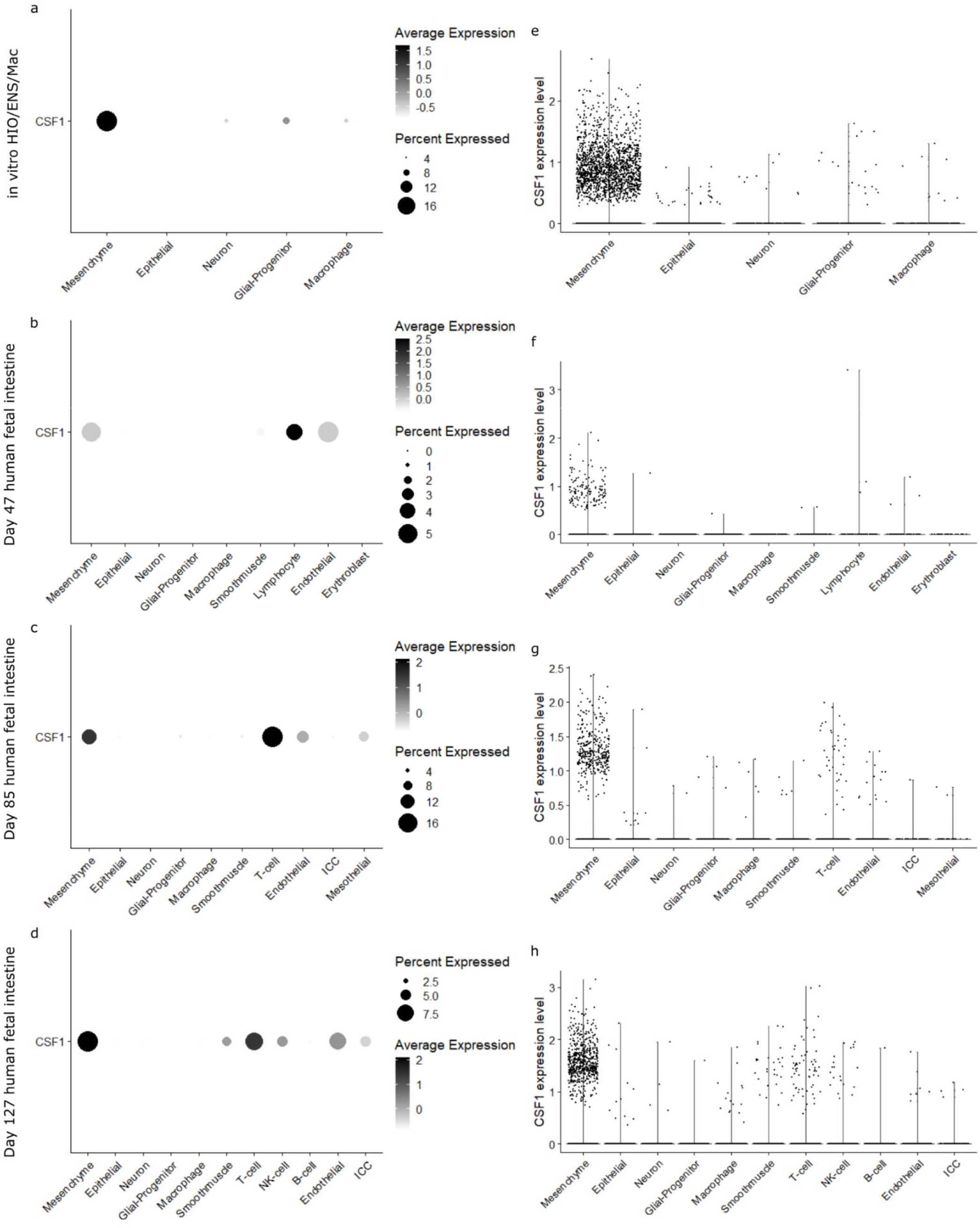
Expression of *CSF1* in HIO/ENS/Mac and fetal intestine scRNAseq datasets. **a-d,** Dot plots of *CSF1* expression in presumptively annotated unsupervised clusters of the day 37 organoid (a), day 47 (b), day 85 (c), day 127 fetal intestine (d) datasets. Percent Expressed, percentage of cells within the cluster expressing the gene. **e-h,** Violin plots of *CSF1* expression of the day 37 organoid (a), day 47 (b), day 85 (c), day 127 fetal intestine (d) datasets. Each dot represents a cell.

**Figure S3.**
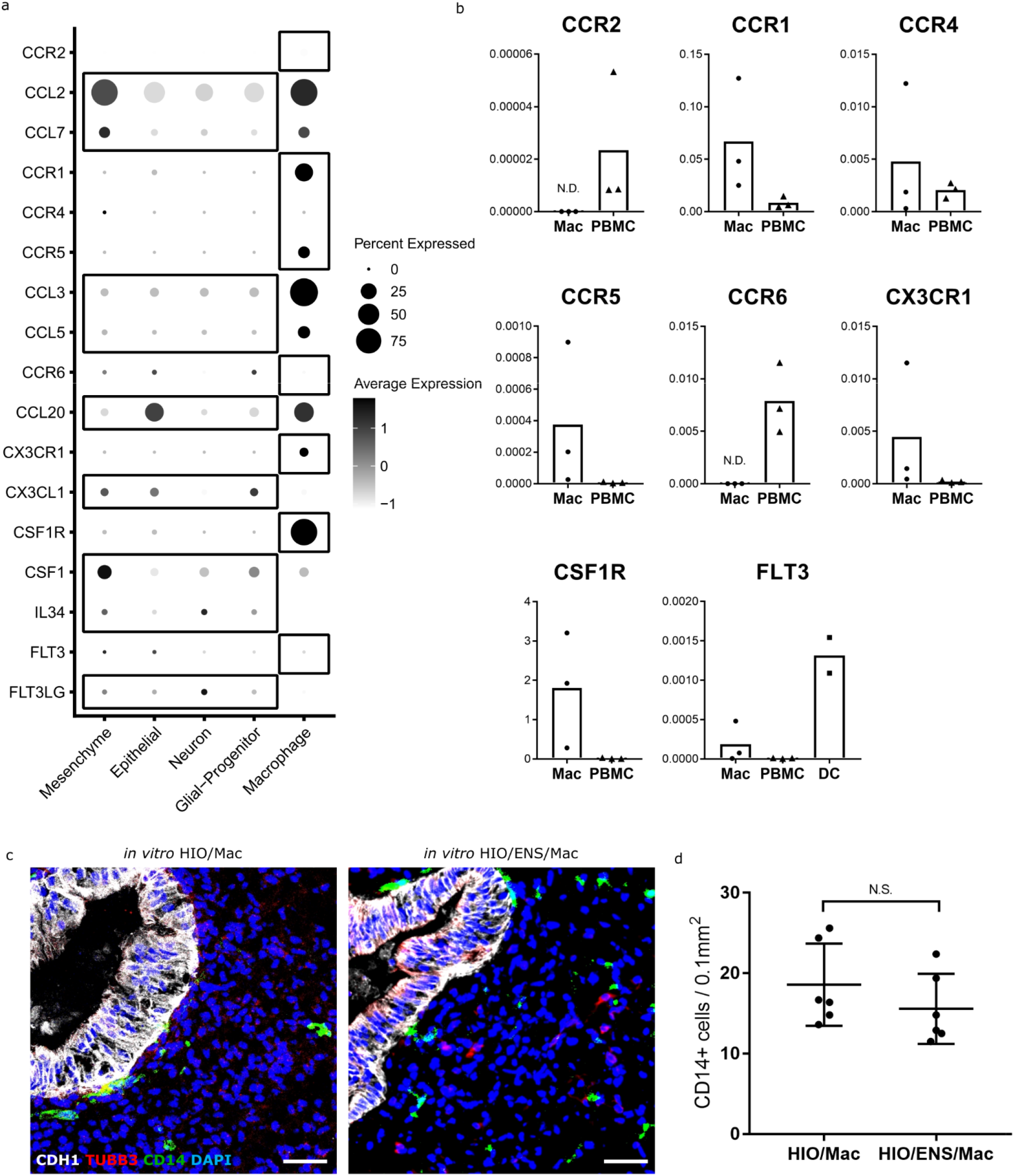
Macrophage recruitment and retention in HIO. **a,** Dot plot of ligand and receptors gene expression involved in macrophage recruitment in HIO/ENS/Mac. **b,** qPCR of receptor genes inquired in (a) in hiPSC-derived macrophages (Mac) prior to co-culture with HIO in comparison to peripheral blood mononuclear cell (PBMC) and dendritic cell (DC). **c,** Immunofluorescence of day 42 HIO co-cultured only with macrophage or with macrophage and vagal neural crest cells (ENS precursor). Epithelium (CDH1), Neuron (TUBB3), macrophage (CD14), nuclei (DAPI). **d,** Quantification of macrophage numbers within the HIOs with or without ENS (c). CD14-positive cells in DAPI-positive area of excluding the CDH1-positive region. Each data point represents an organoid. Mac, n = 3, PBMC, n = 3, DC, n = 2, biological replicates (b). HIO/Mac, n = 6, HIO/ENS/Mac, n = 6 (d). Mean (b). Mean & s.d. (d). p = 0.2996, student’s t-test. Scale bar, 45μm (c).

**Figure S4.**
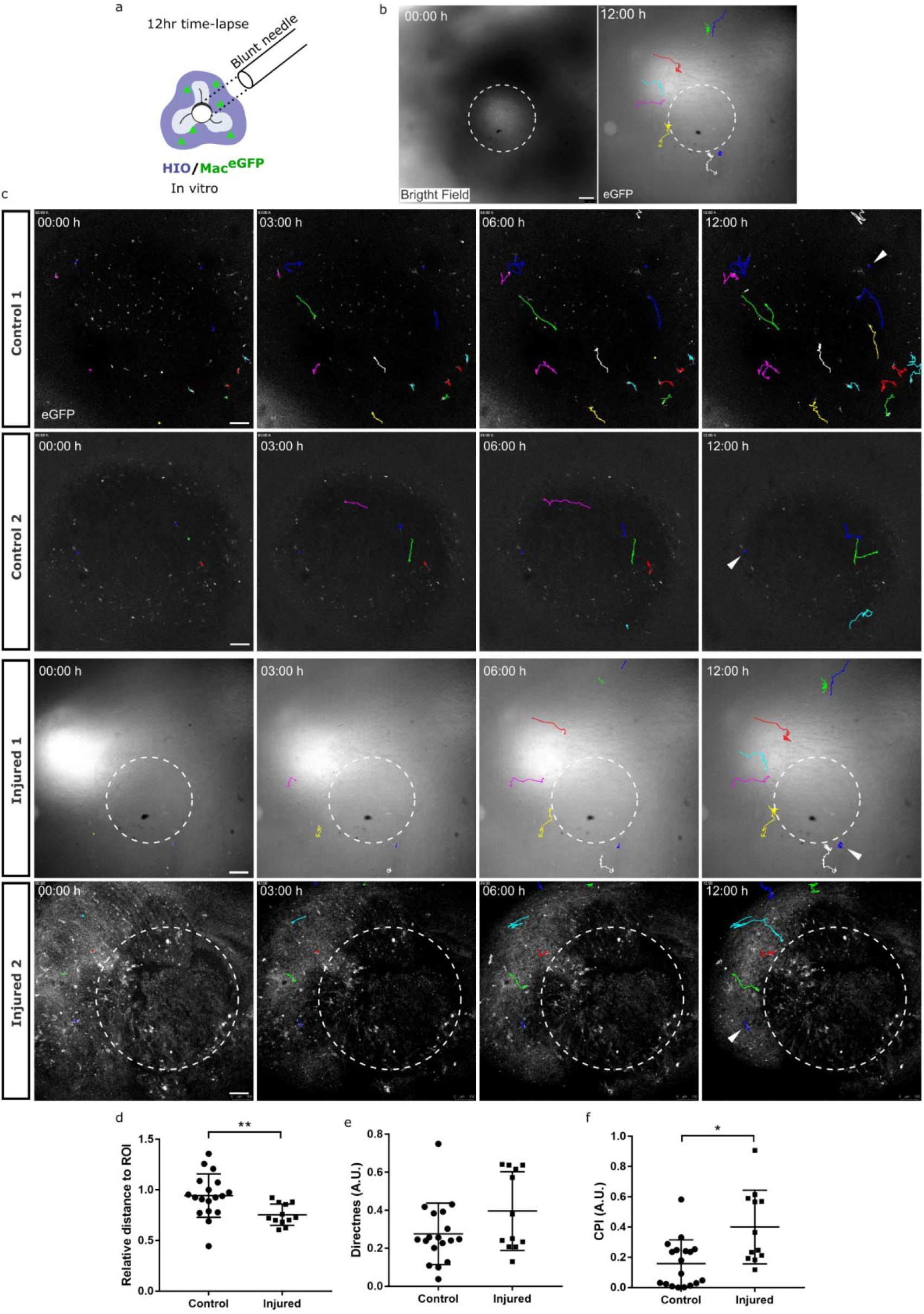
Macrophages in HIO migrate to the wound site upon injury. **a,** Schematic of the *in vitro* organoid injury. **b,** Demarcation of the injury as indicated by the bright field image and the corresponding hiPSC^eGFP^-derived macrophages (Mac^eGFP^) tracing. **c,** Representative GFP fluorescence images from the 12 hour time lapses of control and injured *in vitro* HIO/Mac^eGFP^. Colored lines and dots: tracing of the macrophage location over time. Dotted circle: region of the injury. Arrowhead: stationary points in the organoid to track the drifting of the entire sample. **d,** Relative distance of the macrophages, at 12 hour vs. 0 hour, from the center of the injury (Injured) or an arbitrary point within the image determined blinded (control). **e,** A measure of straightness of cell trajectories quantified as directness. **f,** Degree of directed migration towards a region of interest quantified as chemotactic precision index (CPI). See Methods for calculation. Control, n = 2 organoids, n = 18 cells, Injury, n = 2 organoids, n = 12 cells, each data point represents a cell. Mean & s.d. (d-f). p = 0.0037, Welch’s t-test (d). p = 0.2486 (e), p = 0.0116 (f), Wilcoxon rank sum test (Mann-Whitney). Scale bar, 100μm.

**Figure S5.**
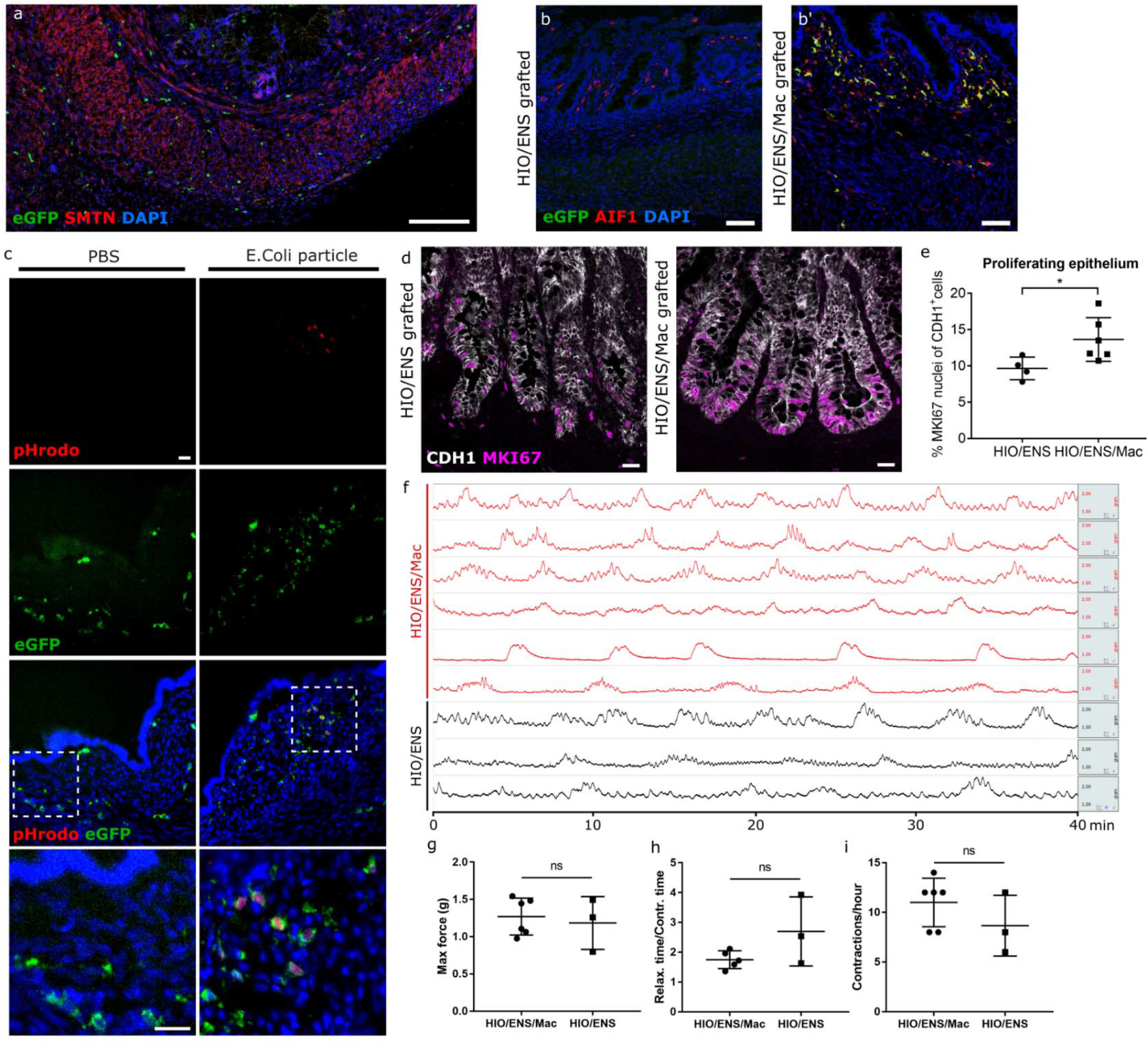
Characterization of grafted organoids. **a,** Representative immunofluorescence image of 10 week grafted HIO/ENS/Mac^eGFP^ probed for eGFP, smooth muscle (SMTN), nuclei (DAPI). **b,b’,** Representative immunofluorescence images of grafted organoids combined without (HIO/ENS) or with hiPSC^eGFP^ derived macrophages (HIO/ENS/Mac^eGFP^). Probed for eGFP and macrophage (AIF1). **c,** Tissue cryosections of grafted HIO/Mac^eGFP^ injected with pHrodo-E.Coli particle conjugates or PBS (vehicle) into the lumen. Tissue sections counterstained with DAPI. Internalized E.Coli signals were only found in flat epithelial areas. **d-e,** Immunofluorescence images of the crypt for proliferating (MKI67) epithelial cells (CDH1) and the quantification of MKI67-positive epithelial cells (e). **f-i,** Isometric force measurements of grafted organoids with or without combined macrophages. Each row represents an individual grafted organoid sample. Maximal force recorded (g), ratio of time required for relaxation to contraction (h), and frequency of contraction (i). See Methods for detail. E.Coli particle injection, n=3; PBS, n=2 grafted organoids (c). HIO/ENS, n = 4, HIO/ENS/Mac, n = 6 (e). HIO/ENS/Mac, n = 6, HIO/ENS, n = 3 (e). Mean & s.d. (e,g-i). p = 0.7303 (g), p = 3252 (h), p = 0.2904 (i), Welch’s t-test. Scale bar, 150μm (a), 50μm (b), 25μm (c), 25μm (d).

**Figure S6.**
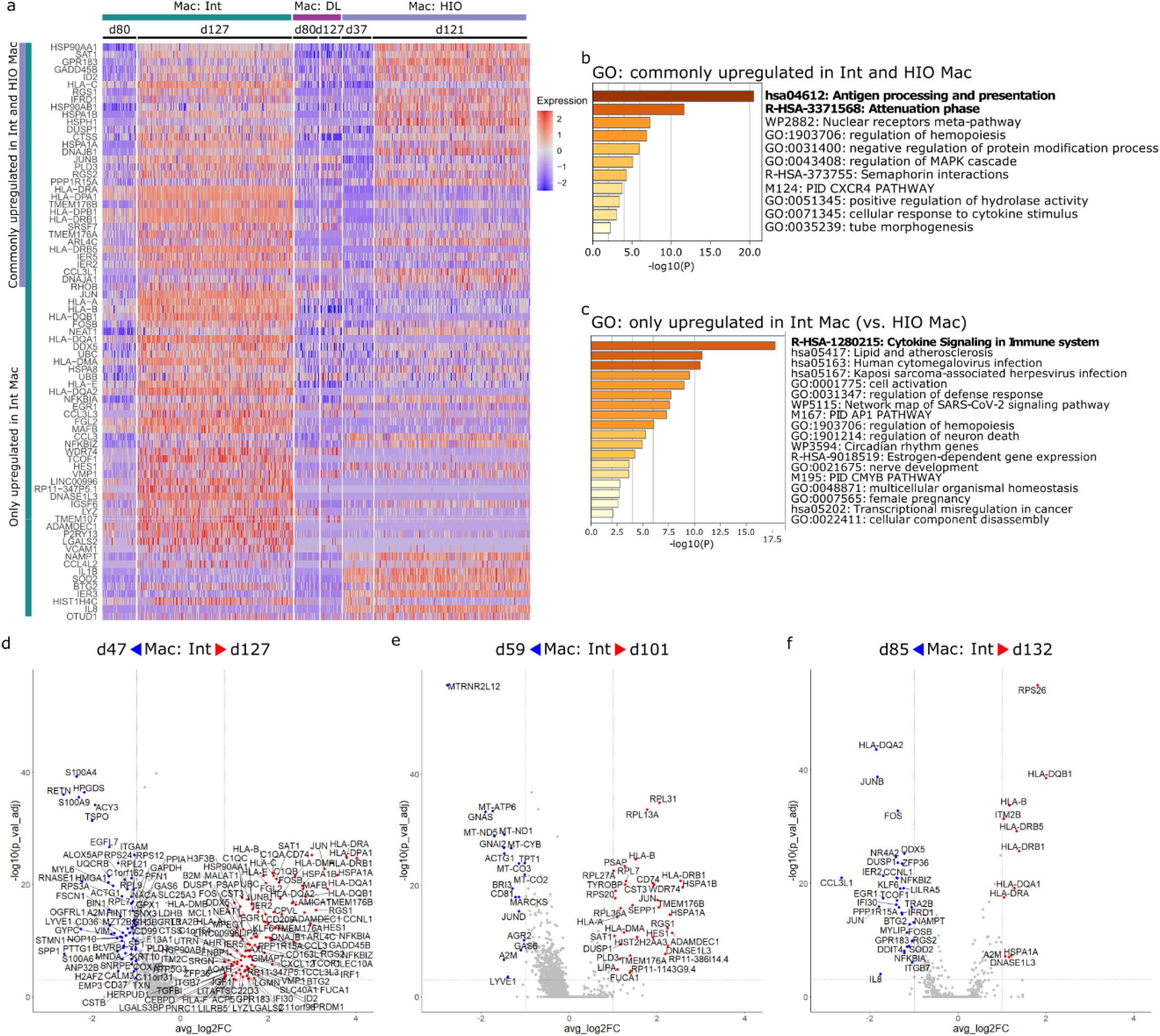
scRNAseq analyses of fetal intestine and organoid macrophages. **a,** Heatmap of all 77 upregulated genes in fetal intestinal macrophages between day 127 and day 80. **b,c,** Gene ontology annotations of differentially expressed genes from (a) for commonly upregulated genes between fetal intestinal and organoid macrophages (b) and upregulated only in fetal intestinal macrophages (c). **d-f,** Volcano plots of differentially expressed genes in fetal intestinal macrophages between day 127 vs. day 47 (d), day 101 vs. day 59 (e), day 132 vs. day 85. Threshold of discovery (dotted line), log2fold change > 1, adjusted p-value < 0.001, Wilcoxon rank sum test (b-f).

**Figure S7.**
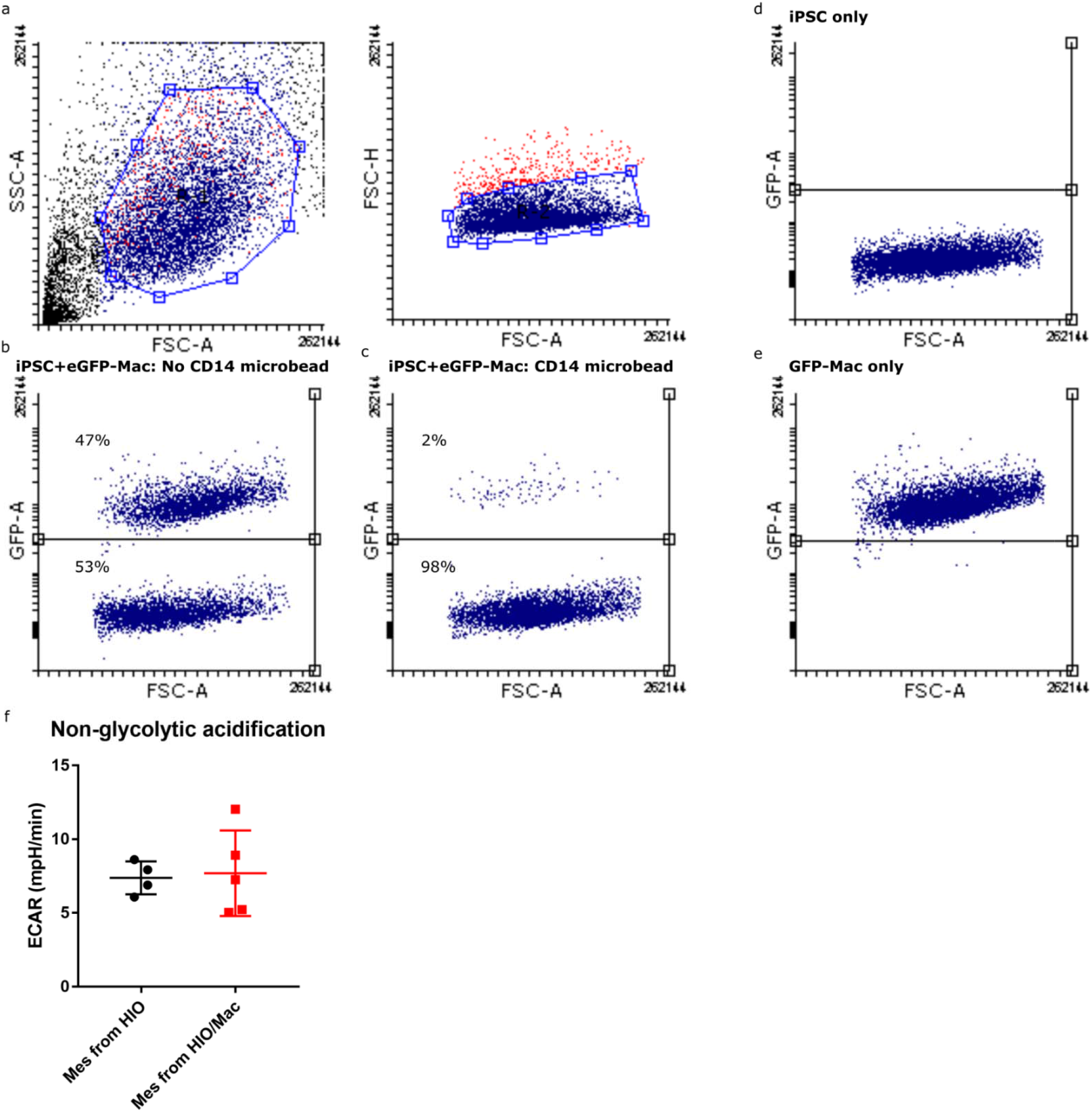
Efficiency of macrophage removal with microbead and ECAR baseline for the glycolytic stress. **a,** Gating for cytometry of 1:1 cell mix of hiPSC^wt^ and hiPSC^eGFP^-derived macrophage. **b,c,** Percent GFP-positive macrophages of the cell mix without (b) or with (c) anti-CD14 microbead mediated cell removal. **d,e,** Percent GFP-positive cells with only hiPSCwt (d) or only hiPSC^eGFP^-derived macrophage. **f,** Extracellular acidification rate (ECAR) of mesenchymal cells (Mes) in the absence of glucose. Mean & s.d, p = 0.8452, student’s t-test (f).

**Figure S8.**
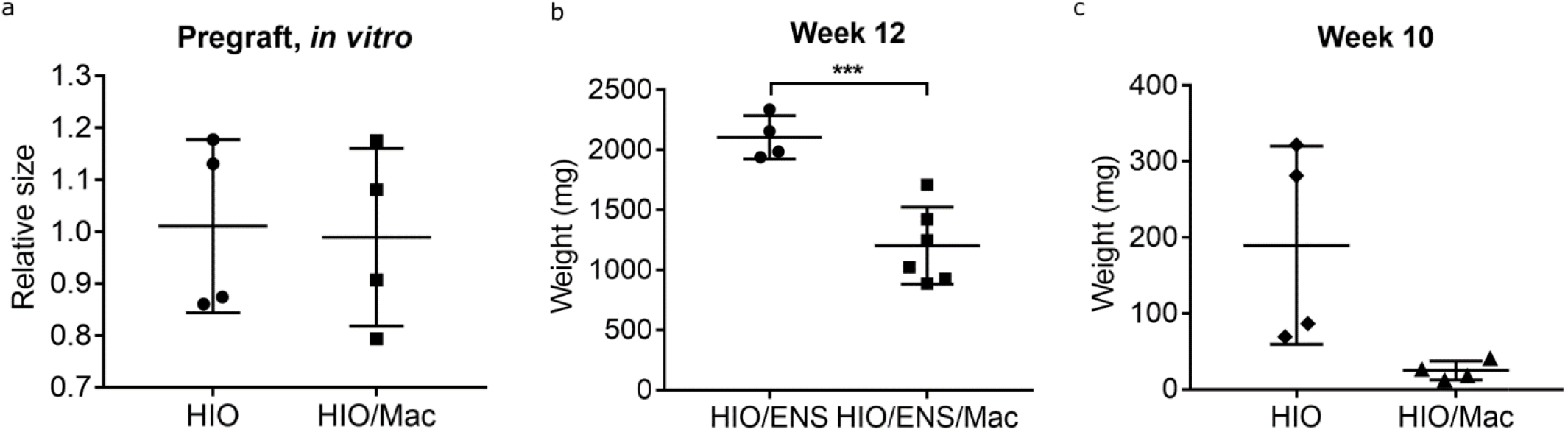
Size and weight of *in vitro* and grafted organoids. **a,** Relative size of HIOs and HIO/Mac *in vitro* shortly after the combination procedure (day 42, two weeks since the start of co-culture) prior to engraftment. **b,c,** Weights of the whole organoids grafted for 12 weeks (b) and 10weeks from (a) (c). HIO, n = 4, HIO/Mac, n = 4 (a,c). HIO/ENS, n = 4, HIO/ENS/Mac, n = 6 (b). Mean & s.d, p = 0.8625 (a), p = 0.0005 (b), p = 0.0848 (c), Welch’s t-test.

**Figure S9.**
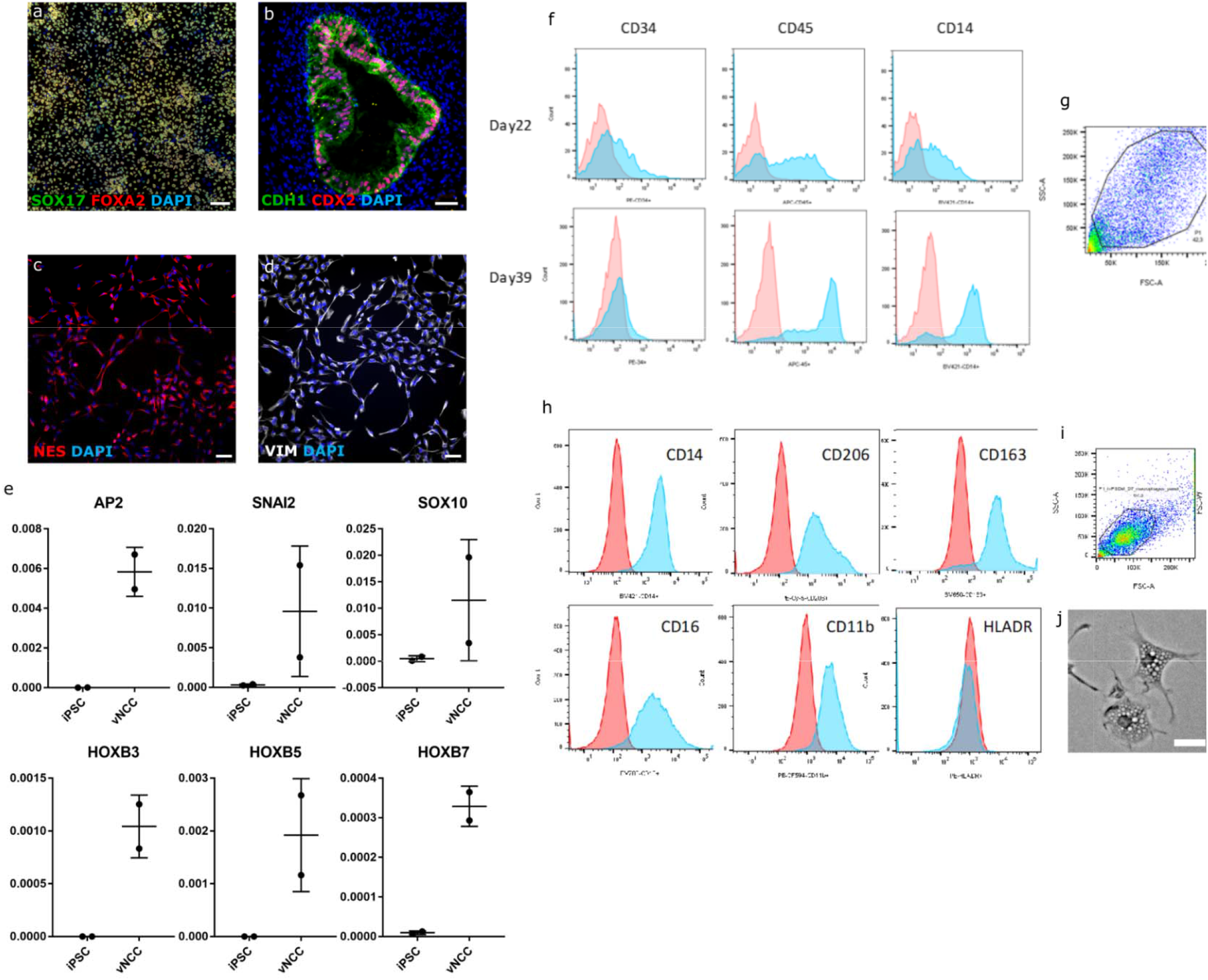
Characterization of hiPSC-derived human intestinal organoid, vagal neural crest, and macrophage. **a,** Immunocytochemistry of markers for definitive endoderm (SOX17, FOXA2) and nucleus (DAPI) of endoderm monolayer during HIO derivation. **b,** Immunofluorescence of intestinal epithelial markers (CDH1, CDX2) in day 28 HIO. **c,d,** Immunocytochemistry for neural crest cell marker (NES, VIM) on the vagal neural crest cells. **e,** qPCR for genetic markers of neural crest (AP2, SNAI2, SOX10) and vagal fate (HOXB3, HOXB5, HOXB7) on the vagal neural crest cells. **f,g,** Flow cytometry histograms showing staining (shaded blues) compared to the compensated unstained control (shaded red) for cell surface markers CD34, CD45 and CD14 on pre-macrophages released from adherent factory embryoid bodies (f-EB) harvested at day 22 and 39 since the beginning of the differentiation (f) and the gating (g). **h,i,** Flow cytometry histograms showing cell surface markers CD14, CD206, CD163, CD16, CD11b and HLADR on macrophages differentiated from pre-macrophages (h) and the gating (i). **j,** Bright field image of the differentiated macrophages. hiPSC, n = 2, VNCC, n = 2 (e) (biological replicate). Mean & s.d. (e). Scale bar, 25μm (j).

